# Plasmacytoid dendritic cell expansion defines a distinct subset of *RUNX1* mutated acute myeloid leukemia

**DOI:** 10.1101/2020.05.11.088872

**Authors:** Wenbin Xiao, Alexander Chan, Michael R. Waarts, Tanmay Mishra, Ying Liu, Sheng F. Cai, Jinjuan Yao, Qi Gao, Robert L. Bowman, Richard Koche, Isabelle S. Csete, Jeeyeon Baik, Sophia Yanis, Christopher Famulare, Minal Patel, Maria E. Arcila, Maximilian Stahl, Raajit K. Rampal, Martin S. Tallman, Yanming Zhang, Ahmet Dogan, Aaron D. Goldberg, Mikhail Roshal, Ross L. Levine

**Affiliations:** Department of Pathology, Hematopathology Service, Memorial Sloan Kettering Cancer Center, New York, NY, USA; Human Oncology & Pathogenesis Program, Molecular Cancer Medicine Service, Memorial Sloan Kettering Cancer Center, New York, NY, USA; Department of Medicine, Leukemia Service, Memorial Sloan Kettering Cancer Center, New York, NY, USA; Department of Pathology, Molecular Diagnostic Laboratory, Memorial Sloan Kettering Cancer Center, New York, NY, USA; Center for Epigenetics Research, Memorial Sloan Kettering Cancer Center, New York, NY, USA; Center for Hematologic Malignancies, Memorial Sloan Kettering Cancer Center, New York, NY, USA; Department of Pathology, Cytogenetics Laboratory, Memorial Sloan Kettering Cancer Center, New York, NY, USA

## Abstract

Plasmacytoid dendritic cells (pDC) are the principal natural type I interferon producing dendritic cells. Neoplastic expansion of pDCs and pDC precursors leads to blastic plasmacytoid dendritic cell neoplasm (BPDCN) and clonal expansion of mature pDCs has been described in chronic myelomonocytic leukemia (CMML). The role of pDC expansion in acute myeloid leukemia (AML) is poorly studied. Here we characterize AML patients with pDC expansion (pDC-AML), which we observe in approximately 5% of AML. pDC-AML often possess cross-lineage antigen expression and have adverse risk stratification with poor outcome. *RUNX1* mutations are the most common somatic alterations in pDC-AML (>70%) and are much more common than in AML without PDC expansion. We demonstrate that pDCs are clonally related to, and originate from, leukemic blasts in pDC-AML. We further demonstrate that leukemic blasts from *RUNX1*-mutated AML upregulate a pDC transcriptional program, poising the cells towards pDC differentiation and expansion. Finally, tagraxofusp, a targeted therapy directed to CD123, reduces leukemic burden and eliminates pDCs in a patient-derived xenograft model. In conclusion, pDC-AML is characterized by a high frequency of *RUNX1* mutations and increased expression of a pDC transcriptional program. CD123 targeting represents a potential treatment approach for pDC-AML.

## Introduction

Plasmacytoid dendritic cells (pDC) are the principal natural type I interferon, interferon-alpha (IFN-α) producing dendritic cells that play critical roles in the immune response(1, 2). Human pDCs can be readily identified by flow cytometry based on high expression of CD123, HLA-DR and CD303/BDCA2 in the absence of other lineage markers(3). pDCs express the cytokine receptor FLT3 (CD135), whose ligand Flt3L is necessary and sufficient for pDC development from both myeloid and lymphoid progenitors(4, 5), suggesting the existence of a robust transcriptional program that drives pDC specification and differentiation(6). TCF4, a master transcription factor(7, 8), acts jointly with MTG16 and additional factors such as BCL11A to promote pDC development (9, 10) by activating transcription factors involved in pDC differentiation *(SPIB, IRF7,* and *IRF8)(7).* IRF family members, particularly IRF7, play a key role in activating IFN-I genes by interacting with other important transcription factors such as RUNX2, SPIB and NFATC3(11–14). CXXC5 and TET2 have been shown to promote *IRF7* expression by maintaining hypomethylation of the *IRF7* promoter in pDCs(15).

The role of pDCs in malignancy has just begun to unfold. The nature of IFN-α producing pDCs has led to the hypothesis that pDCs may possess anti-tumor activities. pDCs via its secreted IFN-α promote NK cell-mediated killing of tumor cells *in vitro* (16–18). Topical treatment with imiquimod, one of the best characterized imidazoquinoline compounds and a TLR7 agonist, leads to the recruitment of pDCs and subsequent secretion of IFN-α and granzyme B, which correlate with regression of basal cell carcinoma and melanoma(19, 20). In contrast, recent studies have shown that pDC accumulation is associated with progression in different human cancer contexts(21–23). Depletion of pDCs inhibits breast cancer cell growth and metastasis in a mouse model(22). It has been shown that tumor associated pDCs possess impaired IFN-α production, resulting in a microenvironment that favors regulatory T-cell expansion. In sum, pDCs may have anti-tumor or pro-tumor effects dependent upon the malignancy and inflammatory context.

Normally, pDCs account for <1% of total nucleated cells in both bone marrow and peripheral blood(24–26). Neoplastic expansion of pDCs or PDC precursors leads to blastic plasmacytoid dendritic cell neoplasm (BPDCN)(27, 28) and clonal expansion of mature pDCs has been reported in chronic myelomonocytic leukemia (CMML)(29). In contrast, the role of pDC expansion in acute myeloid leukemia (AML) remains unelucidated. A few small patient series have reported skin, nodal and marrow pDC expansion in AML (30–36) and a possible correlation with *FLT3^ITD^* mutations has been proposed (37, 38). In 2018, we reported our preliminary analysis of 24 AML patients with pDC expansion suggesting this might represent an entity with distinct genetic and biologic features(39). Here, we performed detailed genetic, transcriptional and functional characterization of AML patients with pDC expansion and investigate potential therapeutic interventions for this specific AML subset.

## Results

### A subset of AML is characterized by increased levels of pDCs

pDCs were identified by their characteristic immunophenotype on flow cytometry (3, 6): specifically low side scatter and CD45 (dim), CD123 (bright), CD303 and HLA-DR surface expression (**Figure 1A-B, Supplemental Figure 1, Supplemental Tables 1 and 2**). The median proportion of pDCs in normal marrow controls was 0.29% (IQR: 0.19-0.37%) (**Figure 1C**). pDCs were markedly depleted in the bone marrow of the majority of AML patients with a median pDC proportion of 0.03% (IQR 0.006-0.14%), a 10-fold reduction compared to normal controls (p<0.001). While none of the normal controls had greater than 1% pDCs, there have been reports of as high as 1.6% pDCs in other normal control cohorts (25). Therefore, 2% was chosen to delineate a stringent cut-off for pDC expansion, which is equivalent to 10 standard deviations (SD) from the mean PDC level of normal controls. Among 850 AML patients, we identified 26 AML patients with ≥2% pDCs in their diagnostic marrow samples. In addition, 16 AML patients had ≥2% pDCs at a later time point after diagnosis. Therefore, 4.9% (42/850) of total AML patients in our study cohort had pDC expansion (hereafter referred to as pDC-AML). In these patients, the proportion of pDC ranged from 2.2% to 35.9%, with a median of 7.7% (IQR 3.4-9.9%, p<0.0001 vs normal control) (**Figure 1C**).

**Figure 1.**
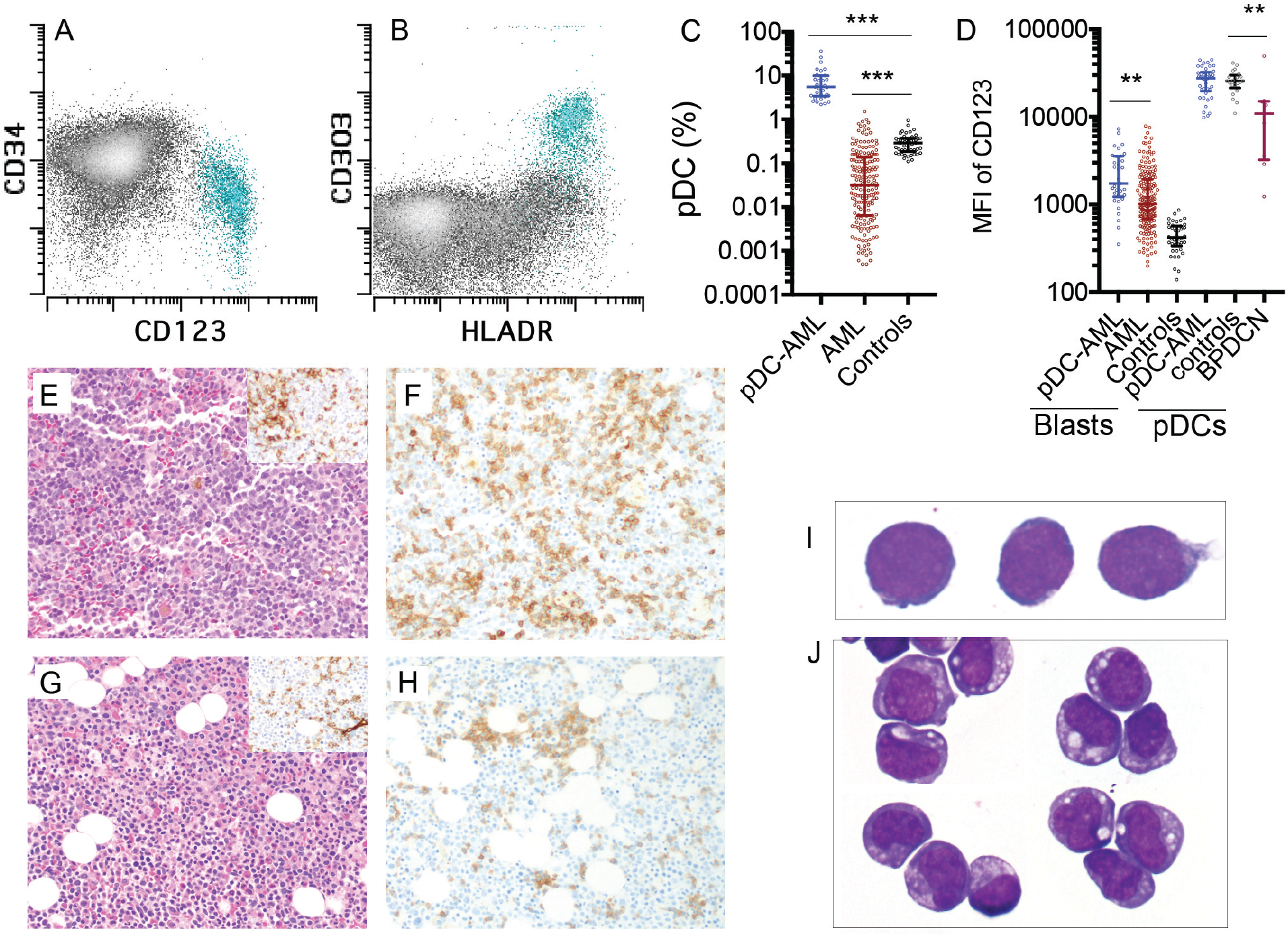
Increased pDCs in a subset of AML. A-B, Flow cytometric identification of pDCs (aqua population represents pDCs). C, pDC proportion (% of WBC) in bone marrow aspirates from AML patients and normal controls (median± IQR). pDC-AML: AML with pDC expansion; AML: AML without pDC expansion; Controls: normal subjects. D, CD123 levels on CD34 positive blasts and pDCs (median± IQR). E, Hematoxylin and eosin stain of bone marrow biopsy from a representative patient with pDC-AML (inset: CD34 immunostain). F, Anti-CD123 immunostain of the patient from panel E. G, Hematoxylin and eosin stain of bone marrow biopsy from another representative patient with pDC-AML (inset: CD34 immunostain). H, CD123 immunostain of the patient from panel G. I-J, Wright-Giemsa stain of flow sorted leukemic blasts (I) and pDCs (J) from pDC-AML. ** p<0.01, *** p<0.001.

### pDCs and blasts from pDC-AML are immunophenotypically distinct to those from AML without pDC expansion

We next performed immunophenotypic characterization of pDCs in pDC-AML samples and compared this to a cohort of BPDCN samples. CD123 was universally expressed on BPDCN cells, albeit at lower levels compared to normal PDCs as previously reported (median of mean fluorescence intensity (MFI): 10855, p=0.0045 vs normal controls, p=0.0015 vs pDC-AML) (**Figure 1D**)(40). By contrast we found that pDCs in pDC-AML patients had high levels of CD123 expression similar to pDCs from normal controls (MFI: 27613 vs 25611, p=0.7). pDC distribution in AML, as demonstrated by CD123 immunohistochemical staining, showed a predominantly interstitial pattern throughout the marrow intermixed with but distinct from CD34 positive leukemic blasts (**Figure 1E-F**). Although small, loose clusters of pDCs were also noted (**Figure 1G-H**), large aggregates as seen in CMML were not observed. Flow cytometry assisted sorting (FACS)-purified leukemic blasts and pDCs displayed distinct blastic and plasmacytoid morphology, respectively (**Figure 1I-J**). The pDCs from pDC-AML patients expressed CD4, CD123 and HLA-DR at levels comparable to normal pDCs. However, they also frequently showed aberrant expression of CD56 (5/29, 17%), CD34 (25/41, 61%), CD5 (5/29, 17%), CD13 (17/41, 41%) and cytoplasmic TdT (4/10, 40%). Other markers expressed by normal pDCs were frequently downregulated in pDCs from pDC-AML patients, including CD303 (7/10, 70%), CD2 (8/29, 28%), and CD33 (11/33, 33%) (**Supplemental Figure 1 and Supplemental Table 2**). This immunophenotype appears intermediate between normal pDCs and BPDCN(40) and shows maturational continuity with the leukemic blasts(36).

The leukemic blasts from 13/41 (32%) pDC-AML patients showed expression of at least two cross-lineage antigens (including CD2, cytoplasmic CD3, CD5, CD7, CD10, CD19, cytoplasmic CD79a) as compared to only 3/100 (3%) AML patients without pDC expansion (p<0.0001, **Table 1**). The most frequent aberrant cross-lineage markers were: CD7 (19/41, 46%), cytoplasmic CD79a (6/19, 32%), CD19 (11/41, 27%), CD2 (6/41, 15%), CD5 (4/41, 10%) and cytoplasmic CD3 (2/24, 8%). 8/41 (20%) patients met the immunophenotypic criteria of mixed phenotype acute leukemia in contrast to only 2/100 (2%) AML patients without pDC expansion (p<0.0001. Table 1, **Supplemental Figure 1 and Supplemental Table 3**). However, these 10 patients with mixed phenotype in both groups carried a diagnosis of secondary AML, which precludes a diagnosis of de novo mixed phenotype acute leukemia(41). After induction therapy, the number of pDCs significantly declined proportionate to the reduction in leukemic blasts and did not significantly rebound (**Supplemental Figure 2A**).

**Table 1.**
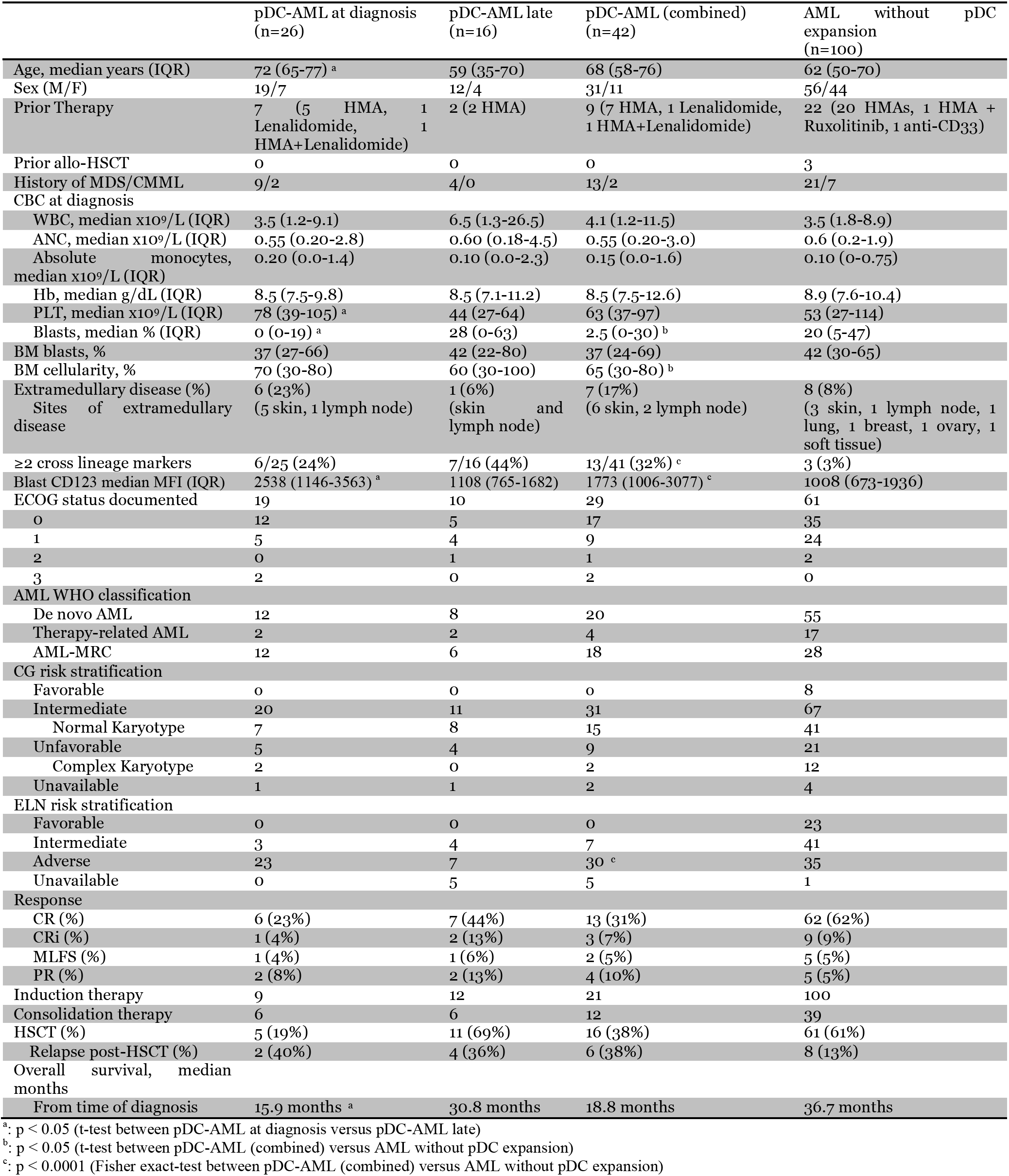
Clinicopathologic features of pDC-AML

### pDC-AML is associated with poor prognosis

pDC-AML patients had a median age of 68 years old (IQR: 58-76) and male predominance (male to female ratio 2.8:1) (**Table 1 and Supplemental Table 4**), similar to AML patients without pDC expansion (referred to as AML). Secondary AML (AML with myelodysplasia-related changes (AML-MRC) and therapy-related AML) were approximately 50% in both cohorts (22/42 in pDC-AML vs 45/100 in AML). Patients with pDC-AML had fewer circulating blasts (median 2.5% vs 20% in AML, p<0.05). Skin involvement was more common in pDC-AML (6/42 (14%) vs 3/100 (3%) in AML, p=0.01). Cytogenetic abnormalities were identified in 25/40 (62.5%, results not available in 2 patients) pDC-AML patients and 9/40 (23%) pDC-AML patients had adverse cytogenetics, not different from the AML cohort (59% and 22%, respectively). The most common abnormality was del7 (5/40, 13%). Only 3/40 (7.5%) had trisomy 13. 30/37 (81%, unable to evaluate in 5 patients) of pDC-AML had adverse risk based on ELN stratification as compared to 35/99 (35%) in the AML cohort (p<0.0001). 21/42 (50%) pDC-AML patients received induction therapy as compared to 100% in the AML cohort (p<0.0001). 6/16 (38%) pDC-AML patients relapsed after allo-HSCT compared to 8/61 (13%) in the AML cohort (p=0.07). The median survival of pDC-AML patients was 18.8 months, which was shorter than the AML cohort (36.7 months, p<0.08) (**Supplemental Figure 2B**). We also compared pDC-AML patients with pDC expansion at diagnosis to those who only manifested expanded pDCs after diagnosis. The former had higher median age (72 vs 59 years old, p<0.05), higher platelet counts (median 78 x10^9^/L vs 44 x10^9^/L, p<0.05) and fewer circulating blasts (0 vs 28%, p<0.05) (**Table 1**). pDC-AML patients with pDC expansion at diagnosis had shorter median overall survival (15.9 vs 30.8 months compared to those with expanded pDCs only after diagnosis, p<0.05) (**Supplemental Figure 2C**).

### pDC-AML is characterized by frequent *RUNX1* mutations

Targeted sequencing analysis revealed that 32/41 (78%) pDC-AML patients had *RUNX1* aberrations, including 29 patients with *RUNX1* mutations (29 patients with 34 total mutations; **Figure 2A**). These mutations were split between missense (18) and nonsense/frame shift (16)with the majority within the RUNT DNA binding domain (**Figure 2B**). 2 patients had atypical *RUNX1* translocations: t(8;21) with *PLAG1* and t(15;21) with *SPATA5L1* as the fusion partners, and one patient had a deletion of the region including *RUNX1.* By comparison, 14/100 (14%) of the AML patients without pDC expansion had *RUNX1* mutations (p<0.0001, Table 1 and data not shown). Other recurrent mutations observed in patients with pDC-AML included *SRSF2* (n=13, 32%), *ASXL1* (n=10, 24%), *TET2* (n=9, 22%), *DNMT3A* (n=7, 17%), *NRAS* (n=7, 17%), *PHF6* (n=6, 15%), *IDH1* (n=4, 10%), *SF3B1* (n=4, 10%), *FLT3* (n=4, 10%) and *TP53* (n=4, 10%). 16/29 (55%) patients with *RUNX1* mutations had at least one mutation commonly observed in clonal hematopoiesis (*TET2*, *ASXL1* and *DNMT3A*). Six patients had *TET2* and *SRSF2* concurrent mutations including the 3 patients with a history of CMML. Based on the variant allele frequencies (VAF) and the blast percentage, *RUNX1* mutations were present in the dominant clones in 26/29 (90%) patients (**Figure 2C and Supplemental Table 5**). 2/4 pDC-AML patients with *TP53* mutations had concurrent *RUNX1* mutations and the VAF of *TP53* mutations in the *RUNX1*-mutant cases was <5%, suggesting a minor *TP53*-mutant subclone and a dominant *RUNX1*-mutant clone.

**Figure 2.**
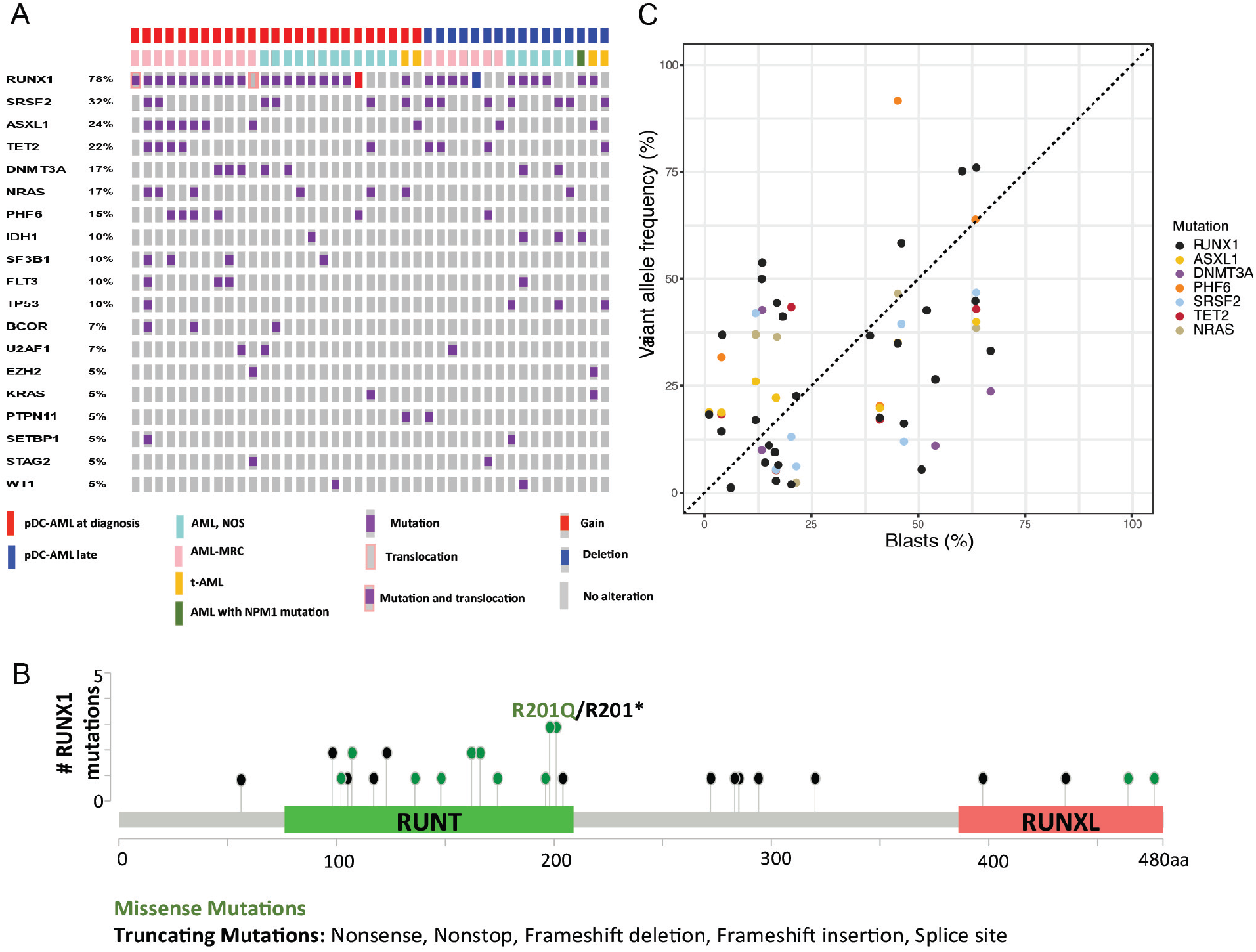
*RUNX1* mutations in pDC-AML. A, Oncoplot of mutations in pDC-AML. B, Lollipop graph of *RUNX1* mutations in pDC-AML. C, VAF of the major mutations as compared to blasts %. AML NOS: AML not otherwise specified; AML-MRC: AML with myelodysplasia related changes; t-AML: therapy related AML.

### pDCs share identical mutations and cytogenetic abnormalities to leukemic blasts

We next assessed whether pDCs in pDC-AML were clonally derived from the leukemic clone. We FACS-purified leukemic blasts and pDCs from 9 patients for paired genomic analysis. Monocytes and T-cells were also sorted from a subset of patients. Targeted sequencing and FISH studies were performed on sorted cells from 7 and 2 patients, respectively. As shown in **Table 2**, the sorted pDCs shared identical mutations and/or cytogenetic abnormalities to the leukemic blasts in all 9 patients. Moreover, the VAF of these mutations were comparable between pDCs and blasts. Monocytes sorted from 3 patients also had the same mutations and abnormalities whereas sorted T cells from 5 patients were negative for all mutations and abnormalities. Collectively, these results indicate that pDCs are neoplastic in origin and are clonally related to the leukemic blasts.

**Table 2.**
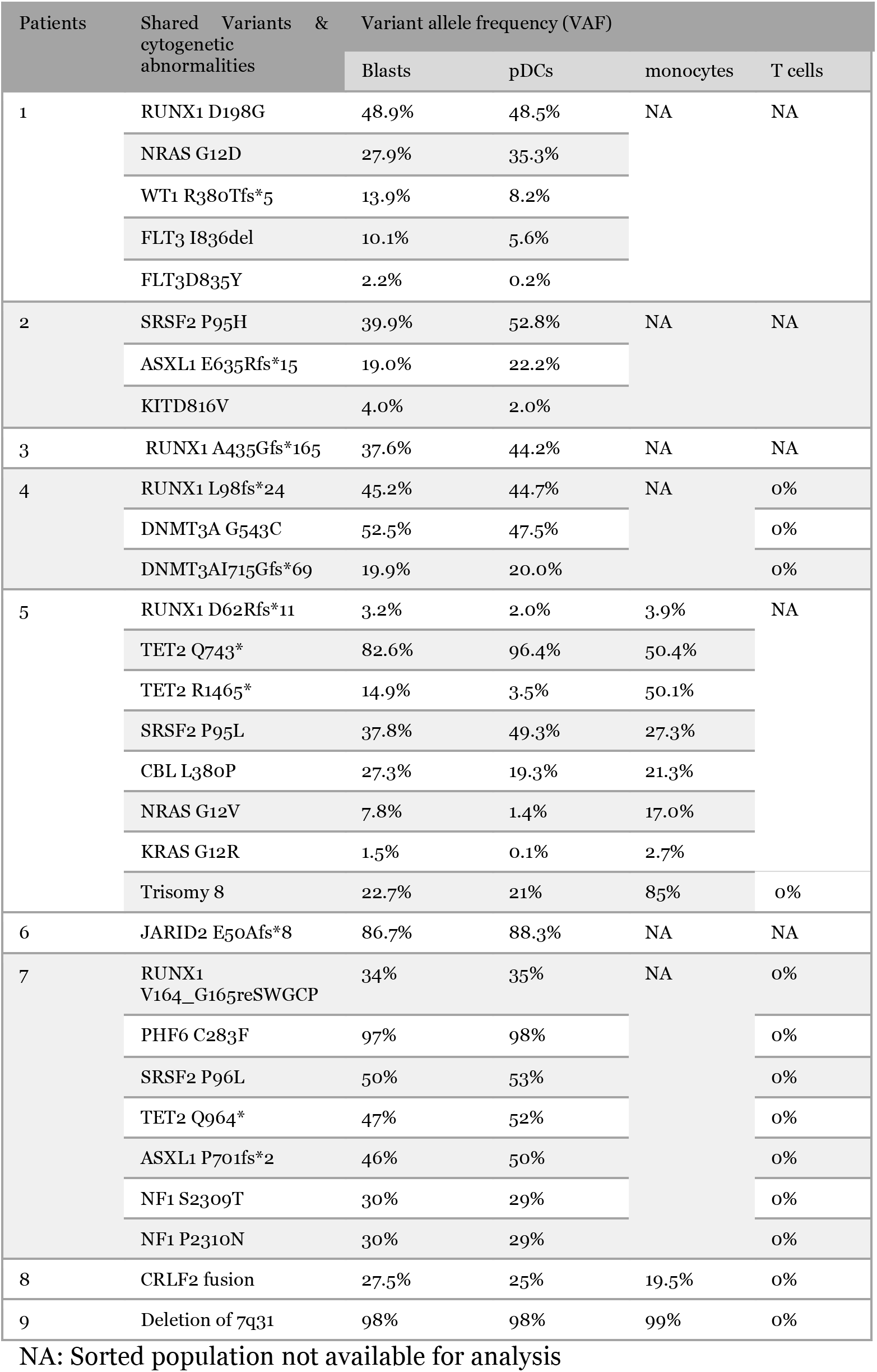
Shared genetic abnormalities between sorted blasts and pDCs from pDC-AML

### Leukemic blasts from pDC-AML show increased pDC differentiation potential

Our genetic data suggest the possibility that pDCs can differentiate from leukemic clones. To address this, we established an *in vitro* culture system that induces pDC differentiation from hematopoietic stem/progenitor cells (HSPC). We cultured cells in FLT3L (100 ng/ml), SCF (10 ng/ml) and TPO (50 ng/ml) containing serum free media (**Supplemental Figure 3**) and assessed pDC differentiation potential with these basal conditions and with the addition of other cytokines/factors. Under these conditions we were able to efficiently differentiate pDCs from cord blood. The pDCs differentiated from cord blood cells showed characteristic surface expression of CD123, HLA-DR and CD303 (**Supplemental Figure 3E-F**) and plasmacytoid cytomorphology (data not shown). Stemregenin 1 (SR1), a small molecule which activates the aryl hydrocarbon receptor, has previously been shown to increase HSC self-renewal(42), platelet production(43) and pDC differentiation(16, 44, 45). Consistent with this, pDC output from cord blood CD34 positive cells was increased by nearly 10-fold by adding SR1 to basal culture conditions **(Supplemental Figure 3A,C)**. pDC output from G-CSF mobilized peripheral blood CD34 positive HSPC was 10-fold lower that that from cord blood HSPC (**Supplemental Figure 3B,D**).

We then FACS-purified leukemic blasts from pDC-AML (n=5), and AML without pDC expansion but with and without *RUNX1* mutations (n=5 and 10, respectively) **(Figure 3A-C)**. We used cord blood CD34 positive cells as controls (n=10). After 2 weeks of *in vitro* culture, cord blood CD34 positive cells generated a mean of 10.8% pDCs (SEM: 1.6%, 95% CI: 7.2%-14.5%). The leukemic blasts from pDC-AML patients had 2.5-fold higher pDC output than cord blood derived cells, with a mean of 26.8% pDCs (SEM 4.6%, 95% CI 14.1%-39.6%, p<0.001 vs cord blood) (**Figure 3A,D**). In contrast, leukemic blasts from AML patients without *RUNX1* mutations and without pDC expansion *in vivo* failed to produce pDCs *in vitro* (**Figure 3C,D**). Interestingly, leukemic blasts from AML patients with *RUNX1* mutations who did not show pDC expansion in their clinical isolates were able to differentiate into pDCs, with a mean of 12.7% pDCs (SEM: 4.6%, 95% CI: 0.4%-25.1%, p<0.04 vs pDC-AML, p<0.001 vs AML without *RUNX1* mutations), equivalent to pDC differentiation/expansion potential from cord blood CD34 positive cells (**Figure 3B,D**).

**Figure 3.**
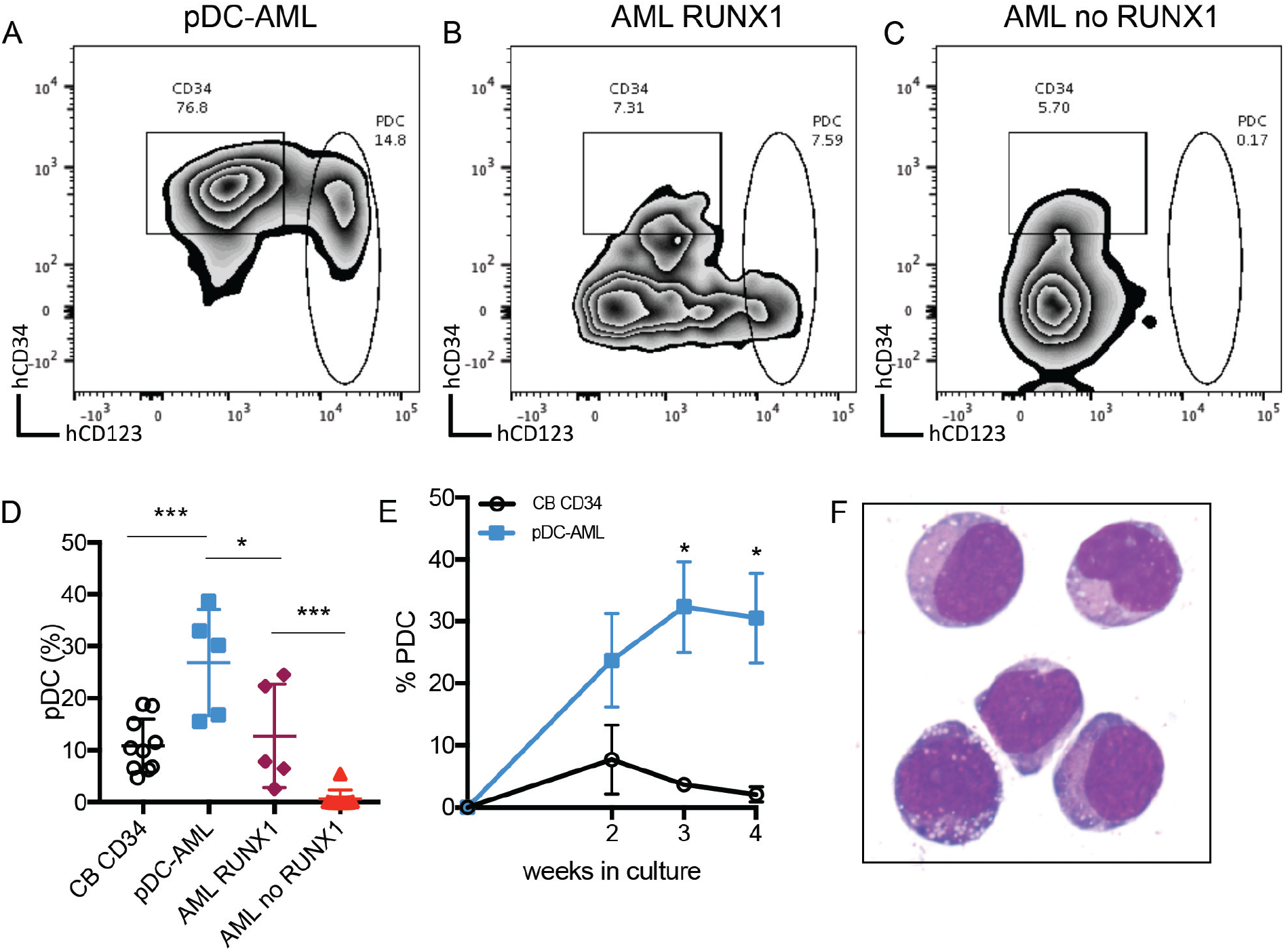
Leukemic blasts from pDC-AML and AML with *RUNX1* mutations have greater differentiation propensity to pDCs *in vitro.* A-C, The sorted leukemic blasts were cultured *in vitro* for 2 weeks and subjected to immunophenotyping by flow cytometry. D, pDC proportion in the culture after 2 weeks was compared. Data are shown as mean±SD. E, Time course experiments showed high pDC proportion persisted from the leukemic blasts of pDC-AML. Data are shown as mean±SEM. F, Wright-Giemsa staining of sorted pDCs from the *in vitro* culture. AML *RUNX1:* AML with *RUNX1* mutations but no pDC expansion; AML no *RUNX1:* AML without *RUNX1* mutations or pDC expansion. * p<0.05, *** p<0.001.

We also studied the dynamic changes of pDC differentiation/expansion *in vitro*. The expanded pDCs generated from cord blood CD34 positive cells peaked at 2 weeks (mean: 7.7%, SEM 5.6%) and gradually declined thereafter (mean 3.7% at 3 weeks and 2.1% at 4 weeks) (**Figure 3E**). Leukemic blasts from pDC-AML had higher pDC output (mean 23.7%, SEM 7.6%) than cord blood CD34 positive cells (p<0.09) at 2 weeks and continued to generate increased levels of pDCs for 4 weeks (mean 32.3% at 3 weeks and 30.5% at 4 weeks, p<0.018 and 0.018, respectively, compared to cord blood) (**Figure 3E**).

pDCs generated *in vitro* had a typical pDC immunophenotype that was CD303 positive and had bright CD123 and HLA-DR expression and were negative for CD11b, CD14, CD19, CD3 or CD56 (data not shown). We also FACS-purified pDCs from *in vitro* cultures at 2 weeks and confirmed cytomorphology was consistent with pDCs (**Figure 3F**). In total, these experiments provide *in vitro* evidence that suggests leukemic blasts from pDC-AML have an intrinsic propensity to differentiate into pDCs.

### Leukemic blasts recapitulate the phenotype of pDC-AML *in vivo*

We next established a patient-derived xenograft (PDX) model to further characterize pDC-AML. PBMC isolated from 6 pDC-AML patients were introduced into sublethally irradiated NSG mice by tail vein injection, and human CD45 positive cells were detected at >0.1% in the peripheral blood of all the recipients at 3 and 6 months (**Supplemental Table 6**)(46). However, only one pDC-AML clinical isolate reproducibly generated overt AML in recipient mice. This PDX showed nearly identical immunophenotype to the original leukemia with high levels of CD34 positive leukemic blasts and increased pDCs (**Supplemental Figure 4**). We then harvested BM cells from primary recipients and transplanted them into secondary NSG recipients in three groups 1)whole bone marrow, 2) FACS purified leukemic blasts, and 3) FACS purified pDCs. Unsorted BM cells (1 million cells per mouse) faithfully recapitulated pDC-AML in secondary recipient mice (**Figure 4**). Sorted leukemic blasts (0.5 million per mouse) also gave rise to AML with increased pDCs (**Figure 4A-D,F**). In contrast, sorted pDCs (0.5 million per mouse) displayed nearly undetectable engraftment in secondary NSG recipient mice up to 8 weeks post transplant (**Figure 4A-B,E**). Collectively, these data provide *in vitro* and *in vivo* evidence indicating that pDCs originate from leukemic blasts but alone cannot initiate leukemia *in vivo*.

### Leukemic blasts from pDC-AML upregulate a pDC transcriptional program

Transcriptional profiling was performed to elucidate the gene regulatory programs underlying the ability of leukemic blasts from pDC-AML to differentiate into pDCs. CD34+ blasts and pDCs were sorted from pDC-AML patients and from normal controls. Based on our identification of *RUNX1* mutations as highly enriched in pDC-AML, we hypothesized that this genetic event may confer a pDC transcriptional program to leukemic blasts. To address this possibility, we sorted CD34+ blasts from AML patients (no pDC expansion) with and without mutations in *RUNX1*. Principal component analysis revealed that pDCs from pDC-AML clustered independently from both normal pDCs and leukemic blasts, consistent with their neoplastic origin and their retention of a pDC transcriptional program **(Figure 5A)**. In order to determine if blasts from pDC-AML or AML with *RUNX1* mutations upregulate a pDC transcriptional signature, we calculated a normalized ssGSEA score based on expression of previously established pDC genes **(Figure 5B, Supplemental Figure 5A)**(47, 48). This analysis showed that blasts from pDC-AML significantly upregulated a pDC transcriptional program as compared to blasts from AML without *RUNX1* mutations (no pDCs). As an orthogonal approach, we identified upregulated genes, with respect to normal marrow CD34+ cells, shared between different groups of leukemic blasts and normal pDCs and confirmed our observations (**Supplemental Figure 5B**). Blasts from AML with *RUNX1* mutation (no pDCs) had greater enrichment of a pDC transcriptional program than blasts from AML without *RUNX1* mutation (no pDCs) but this difference was not statistically significant (**Supplemental Figure 5B**). To identify the specific transcriptional program of blasts from pDC-AML and AML with *RUNX1* mutations, we focused on those upregulated genes shared between these groups and shared with normal pDCs **(Figure 5C, D)**. Gene ontology analysis of this gene subset or of genes upregulated in pDC-AML blasts identified inflammatory pathways and interferon signaling as highly enriched processes, an intriguing finding as these processes contribute to pDC development **(Figure 5E** and **Supplemental Figure 5C**). Consistent with these findings, we observed several genes involved in the interferon response highly upregulated in normal pDCs and neoplastic pDCs, moderately upregulated in blasts from pDC-AML and AML with *RUNX1* mutations, but not upregulated in blasts from AML without *RUNX1* mutations **(Figure 5F)**. In total, this suggests that leukemic blasts from pDC-AML and AML with *RUNX1* mutations may gain an interferon-driven pDC transcriptional program, potentially contributing to the increased pDC output in these settings.

**Figure 4.**
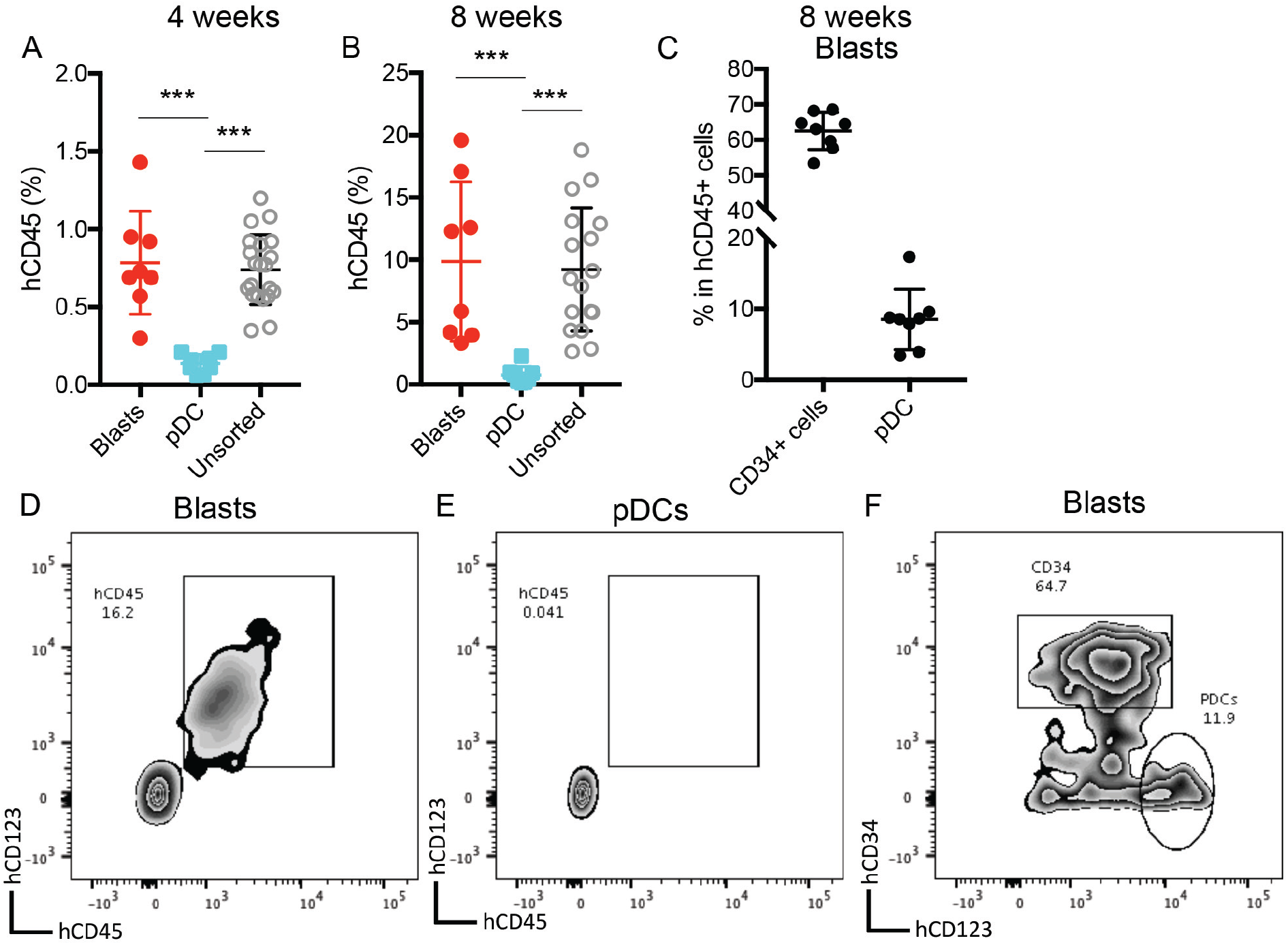
Leukemic blasts differentiate into pDCs *in vivo.* A-B, Bone marrow cells were harvested from the primary NSG mice of pDC-AML leukemic cells as described in supplemental Figure 4 and supplemental Table 6. The unsorted bone marrow cells, purified leukemic blasts and purified pDCs were intravenously injected into secondary NSG mice, respectively. Engraftment of human CD45 positive cells was evaluated from peripheral blood 4 (A) and 8 (B) weeks after transplant. C, pDC-AML phenotype in the secondary NSG mice receiving purified leukemic blasts. Data are shown as mean±SD. D-E, Representative flow plots showing hCD45 positive cells in the secondary NSG mice receiving purified blasts (D) or pDCs (E). F, Representative flow plot showing pDC-AML phenotype in hCD45 positive cells from panel D. ***p<0.001.

**Figure 5.**
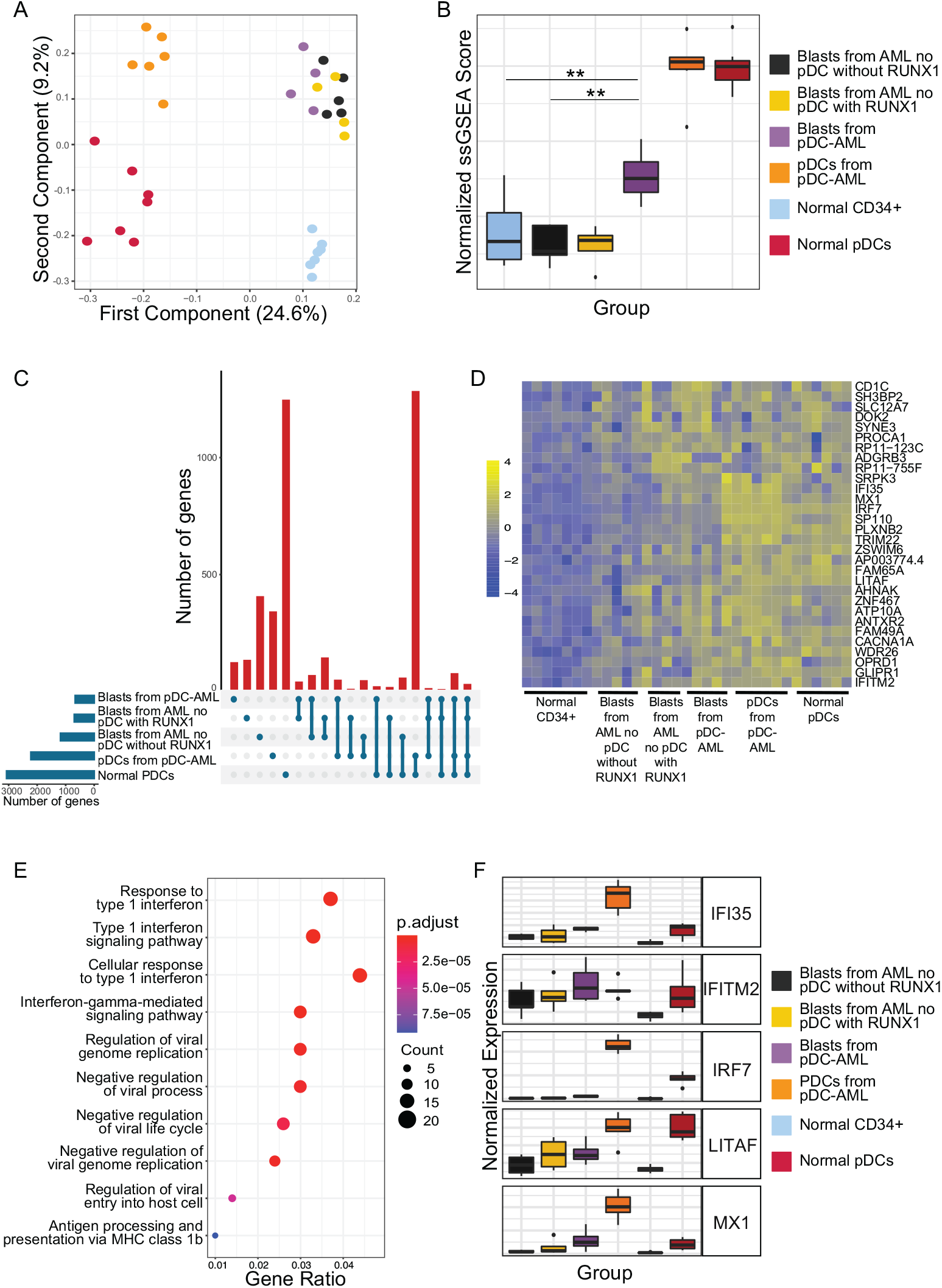
Leukemic blasts from pDC-AML upregulate a pDC transcriptional program. A, Principal component analysis of gene expression in normal marrow CD34+ cells, normal pDCs, blasts from pDC-AML, blasts from AML without *RUNX1* mutations or pDC expansion, and blasts from AML with *RUNX1* mutations but no pDC expansion. B, pDC transcriptional program as evaluated by normalized ssGSEA scores for each group. Scores are calculated based on expression levels of pDC genes. C, Upset plot displaying overlap of upregulated genes between blasts from pDC-AML, blasts from AML without *RUNX1* mutations or pDC expansion, and blasts from AML with *RUNX1* mutations but no pDC expansion, pDCs from pDC-AML, and normal pDCs. All groups are compared to normal marrow CD34+ cells. D, Heatmap showing the 30 upregulated genes shared between blasts from pDC-AML, blasts from AML with *RUNX1* mutations but no pDC expansion, and normal pDCs but not with blasts from AML without *RUNX1* mutations or pDC expansion. E, Gene ontology analysis showing upregulated pathways in blasts from pDC-AML as compared to normal marrow CD34+ cells. F, Expression levels of a subset of interferon-related genes upregulated in pDC-AML. These genes are also among the subset of genes depicted in (D). *p<0.05, **p<0.01.

### Anti-CD123 targeted therapy reduces both pDCs and leukemic blasts in vivo

Tagraxofusp, a fusion protein consisting of interleukin-3 fused to a truncated diphtheria toxin payload with high affinity and avidity for the interleukin-3 receptor-α (CD123), has shown substantive therapeutic efficacy in and therefore is FDA-approved for the treatment of BPDCN(49). Since pDCs are increased in the patients with pDC-AML and the leukemic blasts also expressed higher levels of CD123 compared to AML with no pDC expansion (median of MFI: 1773 vs 1008, p=0.0003) (Figure 1D), we hypothesized that CD123 targeted therapy may have efficacy in pDC-AML. To evaluate tagraxofusp in pDC-AML, unsorted BM cells harvested from primary PDX NSG mice (Supplemental Figure 4 and Supplemental Table 6) were propagated into secondary NSG recipients. After 8 weeks, mice were randomized based on comparable engraftment of hCD45 cells (mean value of hCD45+ cells: 8.4% in PBS cohort, 9.2% in tagraxofusp 0.1mg/kg/d cohort, and 10.0% in tagraxofusp 0.2mg/kg/d cohort, Supplemental Figure 6A). After one cycle of intraperitoneal injection of tagraxofusp (5 days), both 0.1mg/kg/d and 0.2mg/kg/d cohorts showed more than 50% reduction of hCD45+cells (mean: PBS 62.5%, tagraxofusp 0.1mg/kg/d: 33.1%, tagraxofusp 0.2mg/kg/d: 26.8%, p<0.001 PBS vs 0.1mg/kg/d, p<0.001 PBS vs 0.2mg/kg/d) (Figure 6A). Although the proportion of CD34 positive blasts in hCD45 positive population was not significantly affected by the treatment (Figure 6B, D-F), the leukemic burden overall (as evaluated by the proportion of CD34 positive blasts in total white blood cells) was 2-3 fold lower in tagraxofusp treated mice (mean: PBS: 30.7%, tagraxofusp 0.1mg/kg/d: 14.7%, tagraxofusp 0.2mg/kg/d: 11.7%, p<0.0017 PBS vs 0.1mg/kg/d, p<0.0015 PBS vs 0.2mg/kg/d) (Figure 6C). Consistent with the high efficacy of tagraxofusp in BPDCN, pDCs were nearly eliminated by tagraxofusp in both 0.1mg/kg/d and 0.2mg/kg/d cohorts (mean: PBS: 4.8%, 0.1mg/kg/d: 1.3%, 0.2mg/kg/d: 1.4%; p<0.0001 PBS vs 0.1mg/kg/d, or PBS vs 0.2mg/kg/d) (Figure 6B, D-F and Supplemental Figure 6B). Therefore, tagraxofusp is able to effectively eliminate neoplastic pDCs and reduce leukemic burden in an *in vivo* model of pDC-AML.

**Figure 6.**
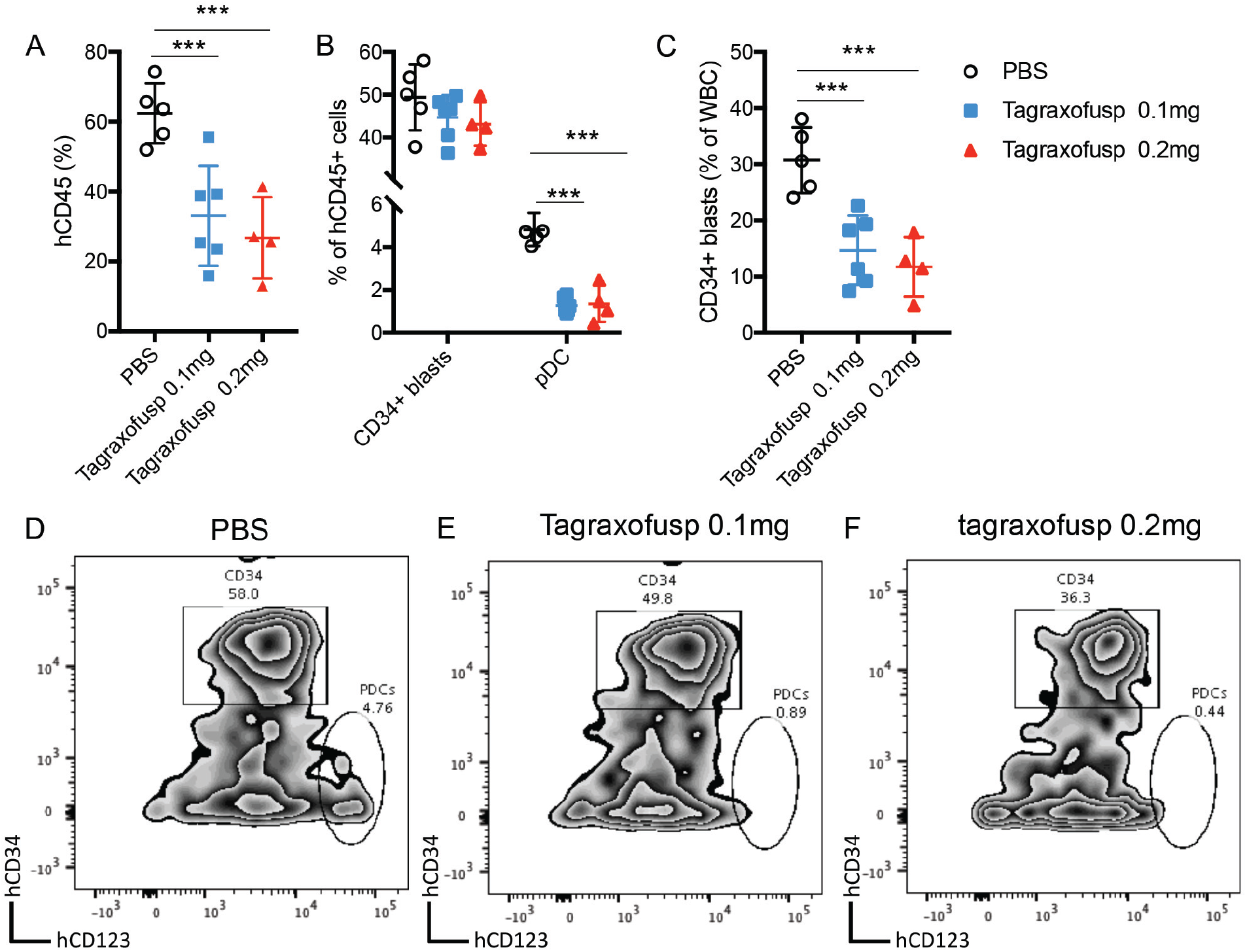
Tagraxofusp treatment results in significant reduction of leukemic blasts and pDCs *in vivo.* A-C, The bone marrow cells harvested from primary NSG mice were intravenously injected into secondary NSG mice. The mice were treated with PBS, tagraxofusp 0.1mg/kg/d and 0.2mg/kg/d for one cycle of intraperitoneal injection. One week after the last dose, hCD45 positive cells were examined in peripheral blood (A). Leukemic blasts and pDC proportions were shown in hCD45 positive compartments (B). CD34 positive leukemic blast proportions in total WBC were also evaluated (C). Data are shown as mean±SD. D-F, Representative flow plots showed the leukemic blasts and pDCs after tagraxofusp treatment (D, PBS, E, tagraxofusp 0.1mg/kg/d, F, tagraxofups 0.2mg/kg/d). *** p<0.001.

## Discussion

Our studies show that a subset of AML patients, most commonly with *RUNX1* mutations, have pDC expansion *in vivo*, pDC differentiation capacity *in vitro* and a distinct transcriptional program suggesting this represents an AML subtype with distinct genetic, transcriptional and biologic features. Moreover, our studies suggest that CD123 targeting may have therapeutic importance in AML with pDC expansion, alone or in combination with other anti-leukemic therapies(50, 51).

The presence of pDC in AML patients raises important diagnostic challenges. Very few patients in our cohort had a history of CMML and the mutational profiles are largely different from that of CMML (including the ones with pDC expansion)(25, 52), arguing against a progression from CMML in the majority of patients. On the other hand, a few patients had greater than 20% pDCs in the marrow with immature pDC morphology, suggesting a potential diagnosis of BPDCN. There are several lines of evidence supporting pDC-AML as an entity distinct from BPDCN. First, skin presentation is relatively infrequent (0nly 14%) in pDC-AML, but is typical (nearly 100%) in BPDCN(41). Second, myeloid blasts are greater than 20% in all patients with pDC-AML. Third, *RUNX1* mutations are present in greater than 70% of pDC-AML but are rare (<5%) in BPDCN(53, 54). Fourth, pDCs from pDC-AML alone are not leukemogenic in contrast to studies showing pDCs from BPDCN can maintain/propagate the diseases(51). These data suggest that pDC-AML is a distinct entity from BPCDN or CMML with pDC expansion, but that there are important shared features which can confound diagnostic evaluation and may inform shared insights into disease pathogenesis and therapy.

Despite the recognition of BPDCN as a distinct entity (29, 41), the role of pDCs in myeloid transformation remains poorly understood. A recent study demonstrated that CMML patients with pDC expansion have higher transformation risk to AML and that pDCs share genetic mutations with the CMML clone(25). However, the role of pDCs in leukemic transformation and maintenance, and in the interaction with malignant cells in different cancer contexts, has not been extensively studied. Our studies suggest that pDCs in AML are derived from the leukemic clone and that there may be interactions between leukemic cells and pDC progeny. Studies have shown that direct contact with E-cadherin and PDL1 expressing myeloma cells could convert pDCs into tumor-promoting cells by suppressing pDC IFN-α production(55, 56). pDCs can also induce regulatory T cells(57), thereby inhibiting anti-tumor response mediated by other effector T cells(58–60). Moreover, leukemia-derived pDCs may have aberrant functionalities, including with respect to cytokine signaling and secretion despite the largely preserved type I IFN program we and others observe(25). Second, pDCs may be a surrogate marker for disease aggressiveness. Since pDCs are markedly depleted in the majority of AML and other high-grade myeloid neoplasms(24, 61), increased pDC differentiation may implicate an early HSC or multipotent progenitor as the cell-of-origin of the myeloid neoplasms, which may correlate with poor outcome(62, 63). Consistent with this hypothesis, cross-lineage marker expression is much more common in pDC-AML than in AML as a whole. Alternatively, pDC differentiation may be indicative of the presence of adverse risk somatic alterations, such as *RUNX1* mutations, which may or may not relate to the cell of origin or differentiation stage of leukemic blasts.

Our data indicate that leukemic blasts from pDC-AML upregulate an interferon-driven pDC transcriptional program, potentially contributing to the increased pDC output. The enrichment of *RUNX1* mutations in pDC-AML implies a crucial role of this genetic event and downstream altered gene regulatory networks in pDC differentiation. Prior studies have shown that pDC-related transcription factors are highly expressed in *RUNX1* mutated AML(64–66), which is further supported by our data. Our study also provides direct evidence that the leukemic blasts from *RUNX1* mutated AML are primed to differentiate into pDCs. *VAV-iCre* inducible *Runx1* deficient mice have markedly decreased Flt3+ lymphoid progenitors including the entire DC compartment(67), consistent with the indispensable role of *Runx1* in lymphoid development(68). Future elucidation of RUNX1 function in DC development requires novel mouse models that have inducible mutant *RUNX1* alleles seen in pDC-AML under the control of pan-hematopoietic versus DC-specific promoters.

Based on the clonal origin and possible pro-leukemic activities of pDCs in AML, we reasoned that targeting pDCs may have therapeutic benefit in pDC-AML. The proof-of-principle experiments demonstrated efficacy of tagraxofusp in eliminating pDCs, consistent with the clinical efficacy in patients with BPDCN(49). Tagraxofusp has previously shown objective responses as a single therapeutic agent in a small subset of AML patients(69). However, this agent did not induce complete responses as monotherapy in our preclinical model of pDC-AML, suggesting that it will be important to combine CD123-targeted therapies with other antileukemic targets to increase therapeutic efficacy in this subset of AML as suggested in tagraxofusp refractory myeloid neoplasms(50, 51). In summary, our study identifies a subset of AML patients with pDC expansion, a high frequency of *RUNX1* mutations, unique transcriptional profile, and increased CD123 expression which may represent a therapeutic target in this high risk AML subset. Future studies are warranted to elucidate the role of pDCs in pDC-AML initiation and maintenance and to develop effective combination therapeutic approaches for this unique AML subset with poor clinical outcome.

## Patients and Methods

### Patients

AML patients were identified from the patient database at the MSKCC between 1/2014 and 12/2019. The key words “plasmacytoid dendritic cells” were searched for flow cytometric and pathology reports. The clinical, morphologic, immunophenotypic and cytogenetic/molecular results were independently re-reviewed by two hematopathologists (WX and MR). A third hematopathologist (AC) also reviewed the data of the patients with increased PDC. A cohort of AML patients without pDC expansion was reported previously(24). A cohort of patients with BPDCN was also identified through similar natural language search. The normal control cohort was previously described with addition of negative marrow specimens (by flow cytometry and morphology) from patients of lymphoma staging(24).

### Human Cells

Umbilical cord blood samples from healthy neonates were obtained from the New York Blood Center under an agreement that includes approval for ethical laboratory research use. Mononuclear cells were separated by density centrifugation using Ficoll-Paque (GE Healthcare) and ammonium chloride red cell lysis. CD34 positive cells were positively selected by magnetic microbeads purchased from Miltenyi. Cryopreserved bone marrow aspirate or peripheral blood mononuclear cell specimens collected at diagnosis were retrieved from our institutional, IRB-approved biospecimen bank.

### Flow cytometry and Cell sorting

For clinical samples, multiparameter flow cytometry was performed on bone marrow aspirates at diagnosis and/or relapse. Briefly, up to 1.5 million cells from freshly drawn bone marrow aspirate were stained with 4-6 10-“color” panels as previously reported, washed, and acquired on a Canto-10 cytometer (BD Biosciences, San Jose, CA)(24). pDC tubes were described in supplemental Table 1. The results were analyzed with custom Woodlist software (generous gift of Wood BL, University of Washington). Cryopreserved live cells were sorted into myeloid blasts, pDCs, monocytes and T cells (as somatic controls) using FACS-Aria Fusion cell sorter(BD Biosciences, San Jose, CA). For cell cultures and PDX samples, similar sample preparation processes were followed. The samples were acquired on a Fortessa cytometry (BD Biosciences, San Jose, CA). The results were analyzed with FlowJo™ Software version 10.6.1 (Ashland, OR: Becton, Dickinson and Company; 2019). The bone marrow cells from PDX mice were sorted using SH800S cell sorter (Sony Biotechnology, San Jose, CA).

### Chromosome and FISH analysis

Conventional chromosome analysis of bone marrow, peripheral blood or fresh tissue was performed following standard procedures with overnight short-term culturing without mitogen. At least 20 metaphase cells were analyzed for a complete chromosome study. The chromosome abnormalities were recorded as per ISCN (2016). In most cases, FISH analysis was performed for recurring chromosome abnormalities in AML and ALL, including t(9;22), MLL, and t(12;21), and in a few cases with inadequate chromosome analysis, extensive FISH tests for t(8;21), inv(16), t(6;9) and IKZF1 were also performed. All FISH probes were from Abbott Molecular (Des Plaines, IL), and its quality and performance were validated in the laboratory.

### Sequencing studies

Bone marrow samples obtained were submitted to a 28-gene amplicon capture-based nextgeneration sequencing (NGS) assay (RainDance) or a larger 400-gene amplicon capture-based next-generation sequencing assay (MSK IMPACT) as previously described. Sorted cells of specific populations from pDC-AML patients were submitted for MSK IMPACT testing. Variants were detected through our clinical workflow.

CD34+ blasts or pDCs were sorted from AML patients or healthy controls into RNA lysis buffer. Total RNA was isolated and RNA integrity and concentration were assessed using a BioAnalyzer. RNA sequencing libraries were generated using SMARTerAmpSeq. Multiplexed libraries were sequenced at the Memorial Sloan Kettering Cancer Center’s core sequencing facility. Approximately 20-30 million paired-end 50 bp reads were sequenced per sample on a Hi-Seq 4000. Fastq files were mapped to the human genome (hg38) and genome wide transcript counting was performed using featureCounts. Differentially expressed genes (DEGs) were identified with DESeq2 and genes with an FDR of 5% were used for downstream analysis. Gene ontology analysis was performed using clusterProfiler. Normalized ssGSEA scores were calculated using normalized expression values and previously published pDC gene sets as input. The algorithm gene set variation analysis (GSVA) was applied(47, 48, 70).

### Cell culture

10,000 FACS-sorted CD34 positive cells were cultured for 14 days in StemSpan serum free culture medium (StemCell Technologies) supplemented with β-mercaptoethanol and penicillin/streptomycin at 37 °C, together with the following cytokines (Peprotech): human fms-like tyrosine kinase 3 ligand (hFLT3L) (100 ng/mL), human stem cell factor (hSCF) (10 ng/mL), human thrombopoietin (hTPO) (50 ng/mL), and Stemregenin 1 (SR1) (1 μg/mL). Half of the medium containing the cytokine cocktail was replaced every 3-4 days.

### Patient-derived xenograft models and tagraxofusp administration

5-6 weeks old, NOD/SCID/IL-2Rg-null female mice were purchased from Taconic and maintained under specific pathogen-free conditions. Mice ages 6-7 weeks were sub-lethally irradiated (2 Gy) up to 24 hr before intravenous (i.v.) injection of various numbers of unsorted human PBMCs, unsorted bone marrow cells or 500,000 sorted leukemic blasts or pDCs. The peripheral blood of reconstituted mice was analyzed every other week after a month. In the treatment experiments, tagraxofusp (gift from Stemline Therapeutics) was diluted in PBS to 10 μg/mL and was administered intraperitoneally (i.p.) at 0.1 mg/kg/d (low dose cohort) or 0.2 mg/kg/d (high dose cohort) for 5 days. Mice were monitored every 1-2 weeks by peripheral blood sampling.

### Statistics

Statistical analyses were performed using Prism software (GraphPad) or DEseq2 for RNA-seq, as indicated. Data are plotted as mean ± SD, except as where indicated in legends. Statistical significance calculations on pairwise comparisons were performed using 2-tailed t tests. Survival analyses were performed on curves generated using the Kaplan-Meier method and groups were compared using the log-rank test. A p value less than 0.05 was considered significant.

### Study approval

This study was approved by the Institutional Review Board at MSKCC and written informed consent was received from participants prior to inclusion in the study. All experiments involving mice were performed in accordance with national and institutional guidelines for animal care and were approved by IACUC at MSK.

## Authorship

Contributions: W.X., M.R., and R.L.L. conceived the study, collected and analyzed the data, and wrote the manuscript. A.C. annotated the clinical and flow data. M.W., R.L.B. and R.K. performed the RNA-seq analysis. T.M., S.F.C., and I.C. performed PDX experiments. Y.L., J.Y., and M.E.A. collected and annotated the mutational data. Q.G. helped cell sorting. J.B., S.Y., C.F. and M.P. coordinated the project. M.S., R.R., M.S.T. and A.D.G. provided clinical data. Y.Z. reviewed cytogenetic data. A.D. analyzed the data. All the authors approved the final version of the manuscript.

## Acknowledgement

This study was supported by the Center for Hematologic Malignancies at MSKCC and in part through the NIH/NCI Cancer Center Support Grant P30 CA008748. M.S. received funding from the MSKCC Clinical Scholars T32 Program under award number 2T32 CA009512-31. This work was supported by National Cancer Institute grants R01CA173636 and R35197594 to RL as well as a Leukemia & Lymphoma Society Specialized Center of Research grant.

## Reference

1. Grouard G, Rissoan M-C, Filgueira L, Durand I, Banchereau J, and Liu Y-J. The Enigmatic Plasmacytoid T Cells Develop into Dendritic Cells with Interleukin (IL)-3 and CD40-Ligand. Journal of Experimental Medicine. 1997;185(6):1101–12.

2. Cella M, Jarrossay D, Facchetti F, Alebardi O, Nakajima H, Lanzavecchia A, and Colonna M. Plasmacytoid monocytes migrate to inflamed lymph nodes and produce large amounts of type I interferon. Nature Medicine. 1999;5(8):919–23.

3. Dzionek A, Fuchs A, Schmidt P, Cremer S, Zysk M, Miltenyi S, Buck DW, and Schmitz J. BDCA-2, BDCA-3, and BDCA-4: three markers for distinct subsets of dendritic cells in human peripheral blood. J Immunol. 2000;165(11):6037–46.

4. Sathe P, Vremec D, Wu L, Corcoran L, and Shortman K. Convergent differentiation: myeloid and lymphoid pathways to murine plasmacytoid dendritic cells. Blood. 2013;121(1):11–9.

5. Shigematsu H, Reizis B, Iwasaki H, Mizuno S-i, Hu D, Traver D, Leder P, Sakaguchi N, and Akashi K. Plasmacytoid Dendritic Cells Activate Lymphoid-Specific Genetic Programs Irrespective of Their Cellular Origin. Immunity. 2004;21(1):43–53.

6. Reizis B. Plasmacytoid Dendritic Cells: Development, Regulation, and Function. Immunity. 2019;50(1):37–50.

7. Cisse B, Caton ML, Lehner M, Maeda T, Scheu S, Locksley R, Holmberg D, Zweier C, den Hollander NS, Kant SG, et al. Transcription factor E2-2 is an essential and specific regulator of plasmacytoid dendritic cell development. Cell. 2008;135(1):37–48.

8. Nagasawa M, Schmidlin H, Hazekamp MG, Schotte R, and Blom B. Development of human plasmacytoid dendritic cells depends on the combined action of the basic helix-loop-helix factor E2-2 and the Ets factor Spi-B. European Journal of Immunology. 2008;38(9):2389–400.

9. Ghosh HS, Ceribelli M, Matos I, Lazarovici A, Bussemaker HJ, Lasorella A, Hiebert SW, Liu K, Staudt LM, and Reizis B. ETO family protein Mtg16 regulates the balance of dendritic cell subsets by repressing Id2. The Journal of experimental medicine. 2014;211(8):1623–35.

10. Wu X, Satpathy AT, Kc W, Liu P, Murphy TL, and Murphy KM. Bcl11a Controls Flt3 Expression in Early Hematopoietic Progenitors and Is Required for pDC Development In Vivo. PloS one. 2013;8(5):e64800.

11. Honda K, Yanai H, Negishi H, Asagiri M, Sato M, Mizutani T, Shimada N, Ohba Y, Takaoka A, Yoshida N, et al. IRF-7 is the master regulator of type-I interferon-dependent immune responses. Nature. 2005;434(7034):772–7.

12. Bao M, Wang Y, Liu Y, Shi P, Lu H, Sha W, Weng L, Hanabuchi S, Qin J, Plumas J, et al. NFATC3 promotes IRF7 transcriptional activity in plasmacy--toid dendritic cells. Journal of Experimental Medicine. 2016;213(11):2383–98.

13. Sasaki I, Hoshino K, Sugiyama T, Yamazaki C, Yano T, Iizuka A, Hemmi H, Tanaka T, Saito M, Sugiyama M, et al. Spi-B is critical for plasmacytoid dendritic cell function and development. Blood. 2012;120(24):4733–43.

14. Chopin M, Preston Simon P, Lun Aaron TL, Tellier J, Smyth Gordon K, Pellegrini M, Belz Gabrielle T, Corcoran Lynn M, Visvader Jane E, Wu L, et al. RUNX2 Mediates Plasmacytoid Dendritic Cell Egress from the Bone Marrow and Controls Viral Immunity. Cell Reports. 2016;15(4):866–78.

15. Ma S, Wan X, Deng Z, Shi L, Hao C, Zhou Z, Zhou C, Fang Y, Liu J, Yang J, et al. Epigenetic regulator CXXC5 recruits DNA demethylase Tet2 to regulate TLR7/9-elicited IFN response in pDCs. Journal of Experimental Medicine. 2017;214(5):1471–91.

16. Díaz-Rodríguez Y, Cordeiro P, Belounis A, Herblot S, and Duval M. In vitro differentiated plasmacytoid dendritic cells as a tool to induce anti-leukemia activity of natural killer cells. Cancer Immunology, Immunotherapy. 2017;66(10):1307–20.

17. Cordeau M, Belounis A, Lelaidier M, Cordeiro P, Sartelet H, Herblot S, and Duval M. Efficient Killing of High Risk Neuroblastoma Using Natural Killer Cells Activated by Plasmacytoid Dendritic Cells. PloSone. 2016;11(10):e0164401.

18. Tel J, Smits EL, Anguille S, Joshi RN, Figdor CG, and de Vries IJ. Human plasmacytoid dendritic cells are equipped with antigen-presenting and tumoricidal capacities. Blood. 2012;120(19):3936–44.

19. Stary G, Bangert C, Tauber M, Strohal R, Kopp T, and Stingl G. Tumoricidal activity of TLR7/8-activated inflammatory dendritic cells. Journal of Experimental Medicine. 2007;204(6):1441–51.

20. Drobits B, Holcmann M, Amberg N, Swiecki M, Grundtner R, Hammer M, Colonna M, and Sibilia M. Imiquimod clears tumors in mice independent of adaptive immunity by converting pDCs into tumor-killing effector cells. The Journal of clinical investigation. 2012;122(2):575–85.

21. Chauhan D, Singh AV, Brahmandam M, Carrasco R, Bandi M, Hideshima T, Bianchi G, Podar K, Tai YT, Mitsiades C, et al. Functional interaction of plasmacytoid dendritic cells with multiple myeloma cells: a therapeutic target. Cancer cell. 2009;16(4):309–23.

22. Sawant A, Hensel JA, Chanda D, Harris BA, Siegal GP, Maheshwari A, and Ponnazhagan S. Depletion of plasmacytoid dendritic cells inhibits tumor growth and prevents bone metastasis of breast cancer cells. J Immunol. 2012;189(9):4258–65.

23. Sisirak V, Faget J, Gobert M, Goutagny N, Vey N, Treilleux I, Renaudineau S, Poyet G, Labidi-Galy SI, Goddard-Leon S, et al. Impaired IFN-alpha production by plasmacytoid dendritic cells favors regulatory T-cell expansion that may contribute to breast cancer progression. Cancer research. 2012;72(20):5188–97.

24. Xiao W, Goldberg AD, Famulare CA, Devlin SM, Nguyen NT, Sim S, Kabel CC, Patel MA, McGovern EM, Patel A, et al. Loss of plasmacytoid dendritic cell differentiation is highly predictive for post-induction measurable residual disease and inferior outcomes in acute myeloid leukemia. Haematologica. 2019;104(7):1378–87.

25. Lucas N, Duchmann M, Rameau P, Noël F, Michea P, Saada V, Kosmider O, Pierron G, Fernandez-Zapico ME, Howard MT, et al. Biology and prognostic impact of clonal plasmacytoid dendritic cells in chronic myelomonocytic leukemia. Leukemia. 2019.

26. Derolf AR, Laane E, Bjorklund E, Saft L, Bjorkholm M, and Porwit A. Dendritic cells in bone marrow at diagnosis and after chemotherapy in adult patients with acute myeloid leukaemia. Scandinavian journal of immunology. 2014;80(6):424–31.

27. Chaperot L, Bendriss N, Manches O, Gressin R, Maynadie M, Trimoreau F, Orfeuvre H, Corront B, Feuillard J, Sotto JJ, et al. Identification of a leukemic counterpart of the plasmacytoid dendritic cells. Blood. 2001;97(10):3210–7.

28. Jacob MC, Chaperot L, Mossuz P, Feuillard J, Valensi F, Leroux D, Bene MC, Bensa JC, Briere F, and Plumas J. CD4(+) CD56(+) lineage negative malignancies: a new entity developed from malignant early plasmacytoid dendritic cells. Haematologica. 2003;88(8):941–55.

29. Vermi W, Facchetti F, Rosati S, Vergoni F, Rossi E, Festa S, Remotti D, Grigolato P, Massarelli G, and Frizzera G. Nodal and extranodal tumor-forming accumulation of plasmacytoid monocytes/interferon-producing cells associated with myeloid disorders. The American journal of surgical pathology. 2004;28(5):585–95.

30. Dargent JL, Delannoy A, Pieron P, Husson B, Debecker C, and Petrella T. Cutaneous accumulation of plasmacytoid dendritic cells associated with acute myeloid leukemia: a rare condition distinct from blastic plasmacytoid dendritic cell neoplasm. Journal of cutaneous pathology. 2011;38(11):893–8.

31. Song HL, Huang WY, Chen YP, and Chang KC. Tumorous proliferations of plasmacytoid dendritic cells and Langerhans cells associated with acute myeloid leukaemia. Histopathology. 2012;61(5):974–83.

32. Dargent JL, Henne S, Pranger D, Balzarini P, Sartenaer D, Bulliard G, Rack K, and Facchetti F. Tumor-forming plasmacytoid dendritic cells associated with myeloid neoplasms. Report of a peculiar case with histopathologic features masquerading as lupus erythematosus. Journal of cutaneous pathology. 2016;43(3):280–6.

33. Wang P, Feng Y, Deng X, Liu S, Qiang X, Gou Y, Li J, Yang W, Peng X, and Zhang X. Tumor-forming plasmacytoid dendritic cells in acute myelocytic leukemia: a report of three cases and literature review. International journal of clinical and experimental pathology. 2017;10(7):7285–91.

34. Wang M, Chen Y-J, Wang L-R, Wang Y-Z, and Lu J. Plasmacytoid Dendritic Cells Proliferation Coexisted with Acute Myeloid Leukemia. Chinese medical journal. 2018;131(15):1866–7.

35. Hamadeh F, Awadallah A, Meyerson HJ, and Beck RC. Flow Cytometry Identifies a Spectrum of Maturation in Myeloid Neoplasms Having Plasmacytoid Dendritic Cell Differentiation. Cytometry B Clin Cytom. 2020;98(1):43–51.

36. Huang Y, Wang Y, Chang Y, Yuan X, Hao L, Shi H, Lai Y, Huang X, and Liu Y. Myeloid Neoplasms with Elevated Plasmacytoid Dendritic Cell Differentiation Reflect the Maturation Process of Dendritic Cells. Cytometry Part A. 2020;97(1):61–9.

37. Rickmann M, Krauter J, Stamer K, Heuser M, Salguero G, Mischak-Weissinger E, Ganser A, and Stripecke R. Elevated frequencies of leukemic myeloid and plasmacytoid dendritic cells in acute myeloid leukemia with the FLT3 internal tandem duplication. Annals of hematology. 2011;90(9):1047–58.

38. Rickmann M, Macke L, Sundarasetty BS, Stamer K, Figueiredo C, Blasczyk R, Heuser M, Krauter J, Ganser A, and Stripecke R. Monitoring dendritic cell and cytokine biomarkers during remission prior to relapse in patients with FLT3-ITD acute myeloid leukemia. Annals of hematology. 2013;92(8):1079–90.

39. Xiao W, Goldberg AD, Famulare C, Baik J, Gao Q, Tallman MS, Zhang Y, Arcila ME, and Roshal M. Acute Myeloid Leukemia with Plasmacytoid Dendritic Cell Differentiation: Predominantly Secondary AML, Enriched for RUNX1 Mutations, Frequent Cross-Lineage Antigen Expression and Poor Prognosis. Blood. 2018;132(Supplement 1):2789–.

40. Wang W, Khoury JD, Miranda RN, Jorgensen JL, Xu J, Loghavi S, Li S, Pemmaraju N, Nguyen T, Medeiros LJ, et al. Immunophenotypic characterization of reactive and neoplastic plasmacytoid dendritic cells permits establishment of a 10-color flow cytometric panel for initial workup and residual disease evaluation of blastic plasmacytoid dendritic cell neoplasm. Haematologica. 2020:haematol.2020.247569.

41. Arber DA, Orazi A, Hasserjian R, Thiele J, Borowitz MJ, Le Beau MM, Bloomfield CD, Cazzola M, and Vardiman JW. The 2016 revision to the World Health Organizationclassification of myeloid neoplasms and acute leukemia. Blood. 2016;127(20):2391–405.

42. Boitano AE, Wang J, Romeo R, Bouchez LC, Parker AE, Sutton SE, Walker JR, Flaveny CA, Perdew GH, Denison MS, et al. Aryl Hydrocarbon Receptor Antagonists Promote the Expansion of Human Hematopoietic Stem Cells. Science (New York, NY). 2010;329(5997):1345–8.

43. Ito Y, Nakamura S, Sugimoto N, Shigemori T, Kato Y, Ohno M, Sakuma S, Ito K, Kumon H, Hirose H, et al. Turbulence Activates Platelet Biogenesis to Enable Clinical Scale Ex Vivo Production. Cell. 2018;174(3):636–48.e18.

44. Thordardottir S, Hangalapura BN, Hutten T, Cossu M, Spanholtz J, Schaap N, Radstake TR, van der Voort R, and Dolstra H. The aryl hydrocarbon receptor antagonist StemRegenin 1 promotes human plasmacytoid and myeloid dendritic cell development from CD34+ hematopoietic progenitor cells. Stem cells and development. 2014;23(9):955–67.

45. Laustsen A, Bak RO, Krapp C, Kjær L, Egedahl JH, Petersen CC, Pillai S, Tang HQ, Uldbjerg N, Porteus M, et al. Interferon priming is essential for human CD34+ cell-derived plasmacytoid dendritic cell maturation and function. Nature communications. 2018;9(1):3525.

46. Wang K, Sanchez-Martin M, Wang X, Knapp KM, Koche R, Vu L, Nahas MK, He J, Hadler M, Stein EM, et al. Patient-derived xenotransplants can recapitulate the genetic driver landscape of acute leukemias. Leukemia. 2017;31(1):151–8.

47. See P, Dutertre C-A, Chen J, Günther P, McGovern N, Irac SE, Gunawan M, Beyer M, Händler K, Duan K, et al. Mapping the human DC lineage through the integration of high-dimensional techniques. Science (New York, NY). 2017;356(6342):eaag3009.

48. Villani AC, Satija R, Reynolds G, Sarkizova S, Shekhar K, Fletcher J, Griesbeck M, Butler A, Zheng S, Lazo S, et al. Single-cell RNA-seq reveals new types of human blood dendritic cells, monocytes, and progenitors. Science (New York, NY). 2017;356(6335).

49. Pemmaraju N, Lane AA, Sweet KL, Stein AS, Vasu S, Blum W, Rizzieri DA, Wang ES, Duvic M, Sloan JM, et al. Tagraxofusp in Blastic Plasmacytoid Dendritic-Cell Neoplasm. New England Journal of Medicine. 2019;380(17):1628–37.

50. Montero J, Stephansky J, Cai T, Griffin GK, Cabal-Hierro L, Togami K, Hogdal LJ, Galinsky I, Morgan EA, Aster JC, et al. Blastic Plasmacytoid Dendritic Cell Neoplasm Is Dependent on BCL2 and Sensitive to Venetoclax. Cancer discovery. 2017;7(2):156–64.

51. Togami K, Pastika T, Stephansky J, Ghandi M, Christie AL, Jones KL, Johnson CA, Lindsay RW, Brooks CL, Letai A, et al. DNA methyltransferase inhibition overcomes diphthamide pathway deficiencies underlying CD123-targeted treatment resistance. The Journal of clinical investigation. 2019;129(11):5005–19.

52. Elena C, Gallì A, Such E, Meggendorfer M, Germing U, Rizzo E, Cervera J, Molteni E, Fasan A, Schuler E, et al. Integrating clinical features and genetic lesions in the risk assessment of patients with chronic myelomonocytic leukemia. Blood. 2016;128(10):1408–17.

53. Menezes J, Acquadro F, Wiseman M, Gomez-Lopez G, Salgado RN, Talavera-Casanas JG, Buno I, Cervera JV, Montes-Moreno S, Hernandez-Rivas JM, et al. Exome sequencing reveals novel and recurrent mutations with clinical impact in blastic plasmacytoid dendritic cell neoplasm. Leukemia. 2014;28(4):823–9.

54. Stenzinger A, Endris V, Pfarr N, Andrulis M, Johrens K, Klauschen F, Siebolts U, Wolf T, Koch PS, Schulz M, et al. Targeted ultra-deep sequencing reveals recurrent and mutually exclusive mutations of cancer genes in blastic plasmacytoid dendritic cell neoplasm. Oncotarget. 2014;5(15):6404–13.

55. Bi E, Li R, Bover LC, Li H, Su P, Ma X, Huang C, Wang Q, Liu L, Yang M, et al. E-cadherin expression on multiple myeloma cells activates tumor-promoting properties in plasmacytoid DCs. The Journal of clinical investigation. 2018;128(11):4821–31.

56. Ray A, Das DS, Song Y, Richardson P, Munshi NC, Chauhan D, and Anderson KC. Targeting PD1-PDL1 immune checkpoint in plasmacytoid dendritic cell interactions with T cells, natural killer cells and multiple myeloma cells. Leukemia. 2015;29(6):1441–4.

57. Lynch JP, Werder RB, Loh Z, Sikder MAA, Curren B, Zhang V, Rogers MJ, Lane K, Simpson J, Mazzone SB, et al. Plasmacytoid dendritic cells protect from viral bronchiolitis and asthma through semaphorin 4a-mediated T reg expansion. The Journal of experimental medicine. 2018;215(2):537–57.

58. Takagi H, Fukaya T, Eizumi K, Sato Y, Sato K, Shibazaki A, Otsuka H, Hijikata A, Watanabe T, Ohara O, et al. Plasmacytoid dendritic cells are crucial for the initiation of inflammation and T cell immunity in vivo. Immunity. 2011;35(6):958–71.

59. Cervantes-Barragan L, Lewis KL, Firner S, Thiel V, Hugues S, Reith W, Ludewig B, and Reizis B. Plasmacytoid dendritic cells control T-cell response to chronic viral infection. Proceedings of the National Academy of Sciences of the United States of America. 2012;109(8):3012–7.

60. Rogers GL, Shirley JL, Zolotukhin I, Kumar SRP, Sherman A, Perrin GQ, Hoffman BE, Srivastava A, Basner-Tschakarjan E, Wallet MA, et al. Plasmacytoid and conventional dendritic cells cooperate in crosspriming AAV capsid-specific CD8(+) T cells. Blood. 2017;129(24):3184–95.

61. Chan A, Zhang Y, Devlin SM, Lewis N, Baik J, Arcila ME, Klimek VM, Roshal M, and Xiao W. Plasmacytoid Dendritic Cell Proportion Is Predictive of Risk and Outcomes in Myelodysplastic Syndromes. Blood. 2019;134(Supplement_1):5439–.

62. Eppert K, Takenaka K, Lechman ER, Waldron L, Nilsson B, van Galen P, Metzeler KH, Poeppl A, Ling V, Beyene J, et al. Stem cell gene expression programs influence clinical outcome in human leukemia. Nature Medicine. 2011;17(9):1086–93.

63. Roshal M, Chien S, Othus M, Wood BL, Fang M, Appelbaum FR, Estey EH, Papayannopoulou T, and Becker PS. The proportion of CD34+CD38low or neg myeloblasts, but not side population frequency, predicts initial response to induction therapy in patients with newly diagnosed acute myeloid leukemia. Leukemia. 2013;27(3):728–31.

64. Simon L, Lavallee VP, Bordeleau ME, Krosl J, Baccelli I, Boucher G, Lehnertz B, Chagraoui J, MacRae T, Ruel R, et al. Chemogenomic Landscape of RUNX1-mutated AML Reveals Importance of RUNX1 Allele Dosage in Genetics and Glucocorticoid Sensitivity. Clinical cancer research: an official journal of the American Association for Cancer Research. 2017;23(22):6969–81.

65. Gerritsen M, Yi G, Tijchon E, Kuster J, Schuringa JJ, Martens JHA, and Vellenga E. RUNX1 mutations enhance self-renewal and block granulocytic differentiation in human in vitro models and primary AMLs. Blood Adv. 2019;3(3):320–32.

66. In ‘t Hout FEM, Gerritsen M, Bullinger L, van der Reijden BA, Huls G, Vellenga E, and Jansen JH. Transcription factor 4 (TCF4) expression predicts clinical outcome in RUNX1 mutated and translocated acute myeloid leukemia. Haematologica. 2019.

67. Satpathy AT, Briseno CG, Cai X, Michael DG, Chou C, Hsiung S, Bhattacharya D, Speck NA, and Egawa T. Runx1 and Cbfbeta regulate the development of Flt3+ dendritic cell progenitors and restrict myeloproliferative disorder. Blood. 2014;123(19):2968–77.

68. Ichikawa M, Asai T, Saito T, Seo S, Yamazaki I, Yamagata T, Mitani K, Chiba S, Ogawa S, Kurokawa M, et al. AML-1 is required for megakaryocytic maturation and lymphocytic differentiation, but not for maintenance of hematopoietic stem cells in adult hematopoiesis. Nature medicine. 2004;10(3):299–304.

69. Frankel A, Liu J-S, Rizzieri D, and Hogge D. Phase I clinical study of diphtheria toxin-interleukin 3 fusion protein in patients with acute myeloid leukemia and myelodysplasia. Leukemia & lymphoma. 2008;49(3):543–53.

70. Hänzelmann S, Castelo R, and Guinney J. GSVA: gene set variation analysis for microarray and RNA-Seq data. BMCBioinformatics. 2013;14(1):7.

