## supplemental Table 1 for "Plasmacytoid dendritic cell expansion defines a distinct subset of *RUNX1* mutated acute myeloid leukemia"

Supplemental Figure 1
Supplemental Figure 2
Supplemental Figure 3
Supplemental Figure 4
Supplemental Figure 5
Supplemental Figure 6
Supplemental Table 1
Supplemental Table 2
Supplemental Table 3
Supplemental Table 4
Supplemental Table 5
Supplemental Table 6

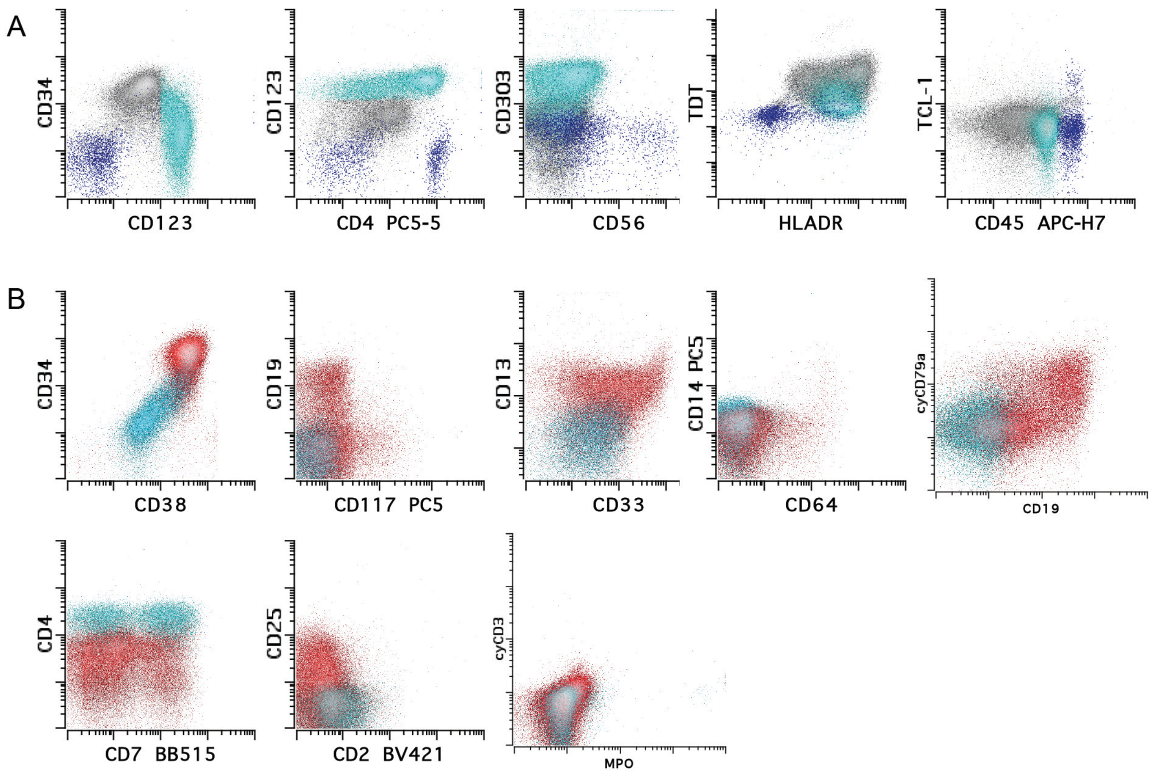


Supplemental Figure 1. Detailed immunophenotype of pDCs and leukemic blasts from a representative pDC-AML patient. A. Flow plots of DC tubes (aqua, pDCs; grey, blasts; blue, lymphocytes). B. Flow plots of myeloid and intracytoplasmic tubes (aqua, pDCs; red, blasts). The pDCs are positive for CD123, CD4, HAL-DR, dim CD45, dim CD34, dim CD38, CD7 (subset) and dim CD33, but negative for CD56, TCL-1, TdT, CD117, CD2, CD5(not shown), CD8 (not shown), CD25, CD13, CD14, CD64, CD19, cyCD79a, cyCD3 and MPO. The blasts are positive for CD34, CD38, HLA-DR, CD117 (minor subset), intermediate CD123, intermediate CD4, dim TdT, CD13, CD33, CD7 (subset), CD25 (subset), CD19 (subset) and cyCD79 (subset), but negative for CD303, CD56, CD14, CD2, cyCD3, MPO and TCL-1. The blasts show myeloid and B lineage mixed phenotype.

Fig S2

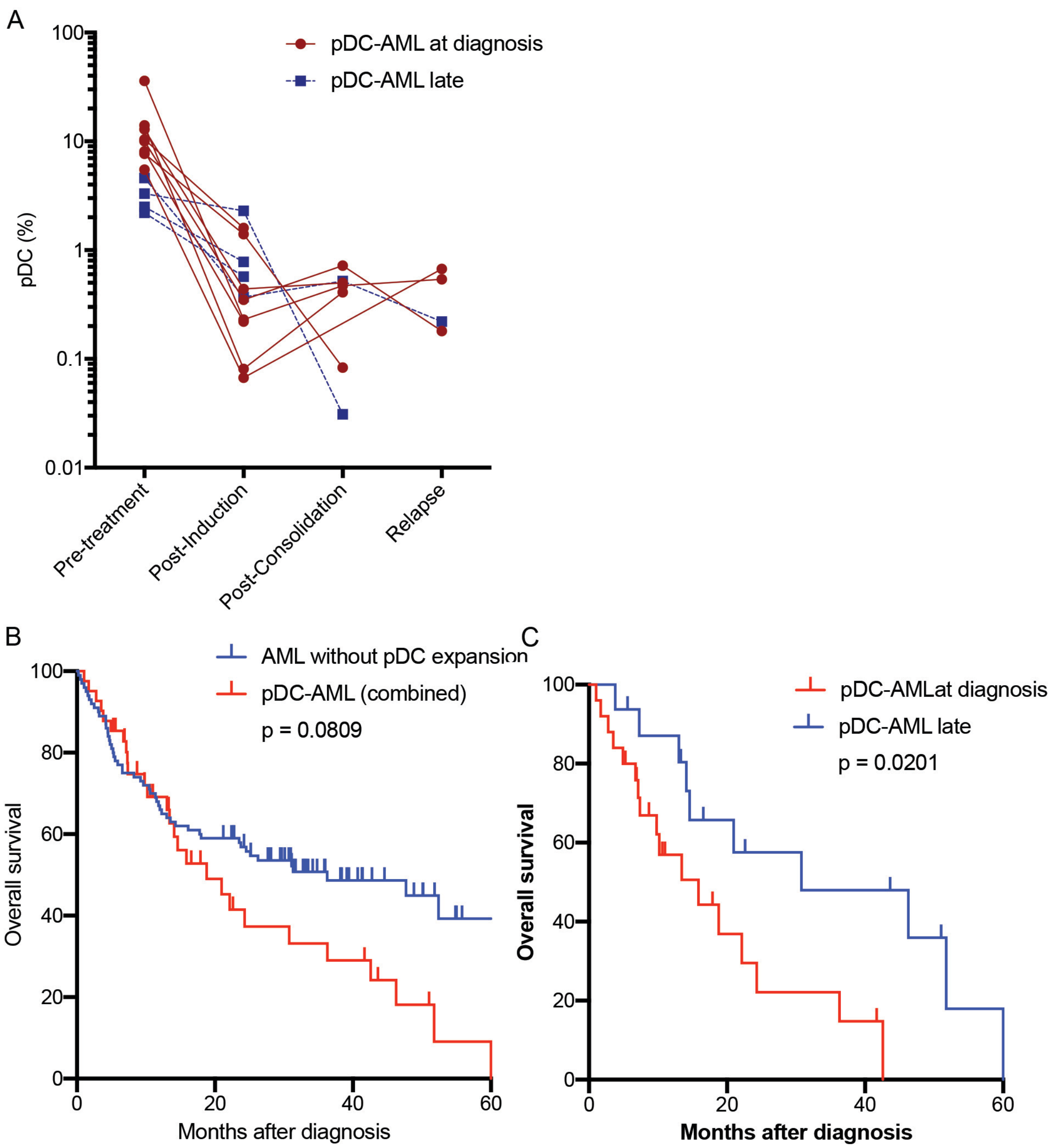


Supplemental Figure 2. Dynamics of pDC after treatment (A) and Kaplan-Meier survival curve (B and C).

Fig S3

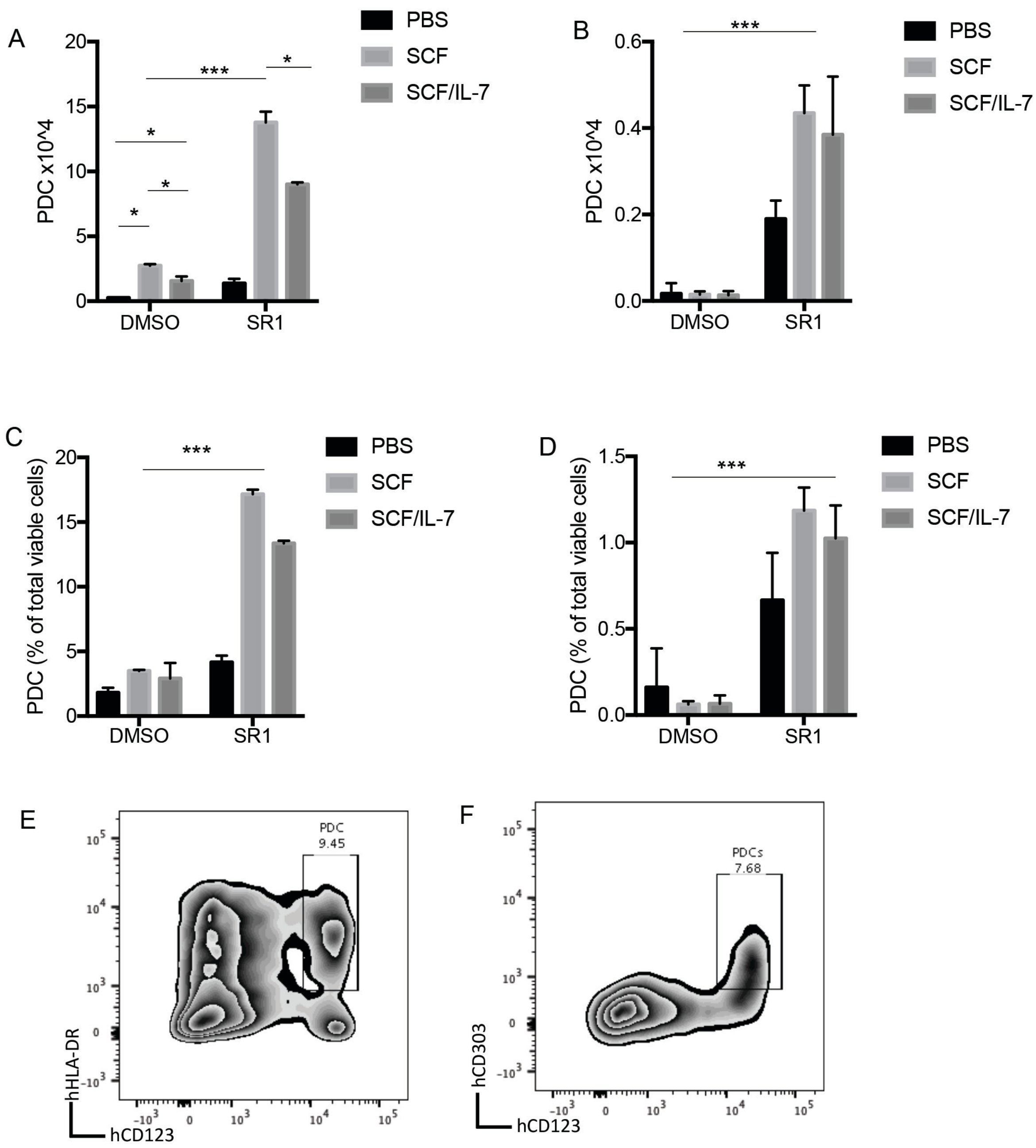


Supplemental Figure 3. pDC differentiation facilitated by SR1. A-B, Purified CD34 positive cells from cord blood (A and C) and G-CSF mobilized peripheral blood mononuclear cells (B and D) were cultured in STEMSPAN serum free medium with hFLT3L 100ng/ml, hTPO 50 ng/ml and cytokines or SR1 as indicated for 2 weeks. E-F, Cells cultured in vitro for 2 weeks (from panel A) with addition of SR1 and SCF 10 ng/ml were immunophenotyped by flow cytometry. pDCs were identified as DAPI-CD3-CD19-CD56-CD14-FceRI-CD123+HLA-DR+CD303+ population.

Fig S4

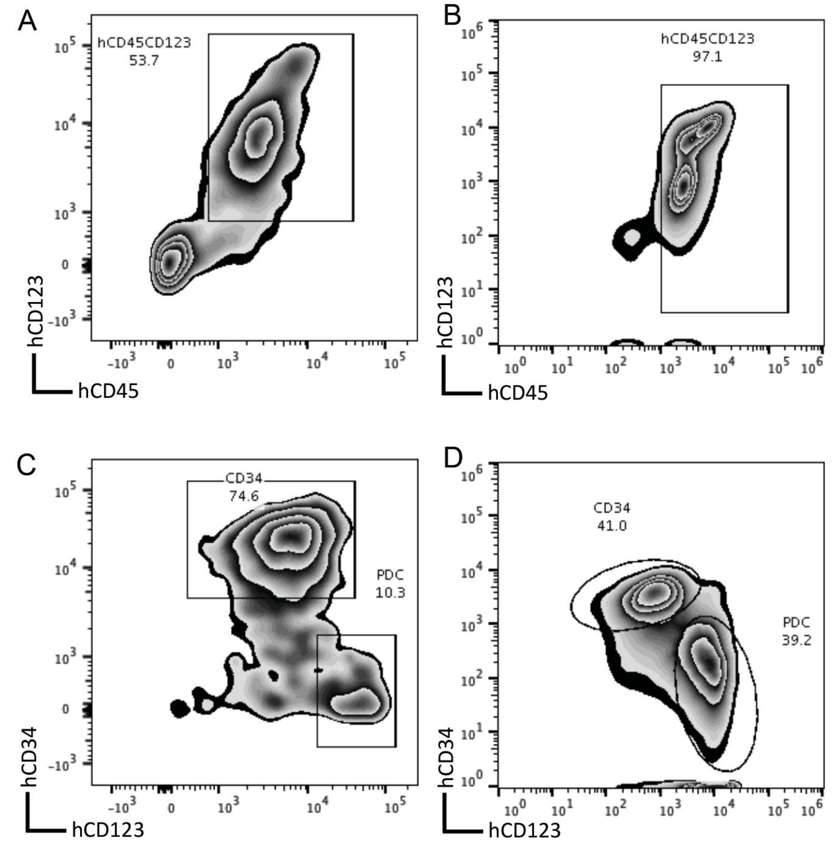


Supplemental Figure 4. Immunophenotype of primary NSG mice mice receiving pDC-AML cells. A and B, hCD45 engraftment in peripheral blood (A) and bone marrow (B) 6 months after transplant. C and D, Leukemic blasts and pDCs in hCD45 positive cells from peripheral blood (C) and bone marrow (D).

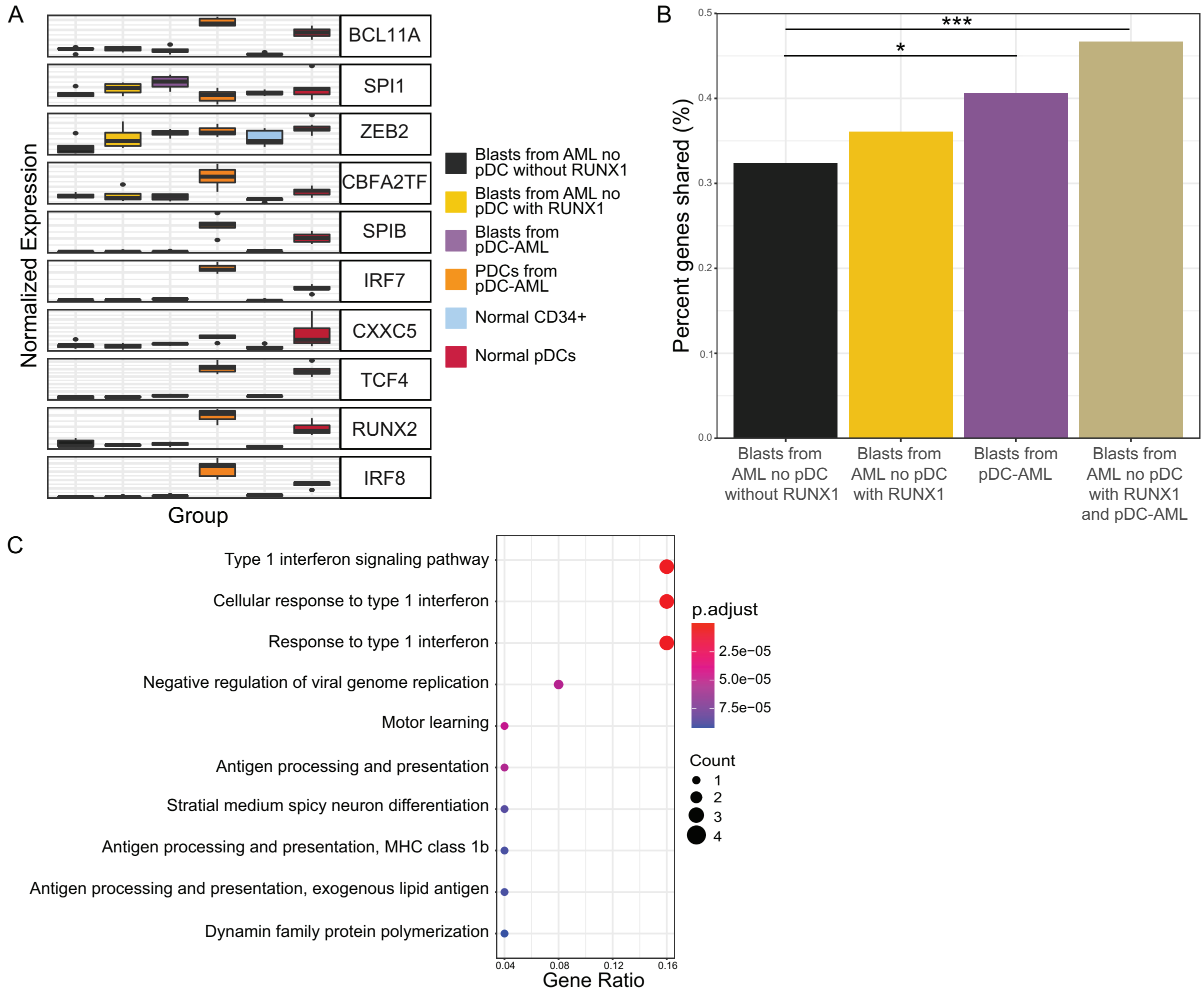


Supplemental Figure 5. A, Expression levels of key pDC genes. B, Shared pDC transcriptional program for each group. C, Gene ontology analysis showing upregulated pathways in the subset of 30 genes from Figure 5C.

Fig S6

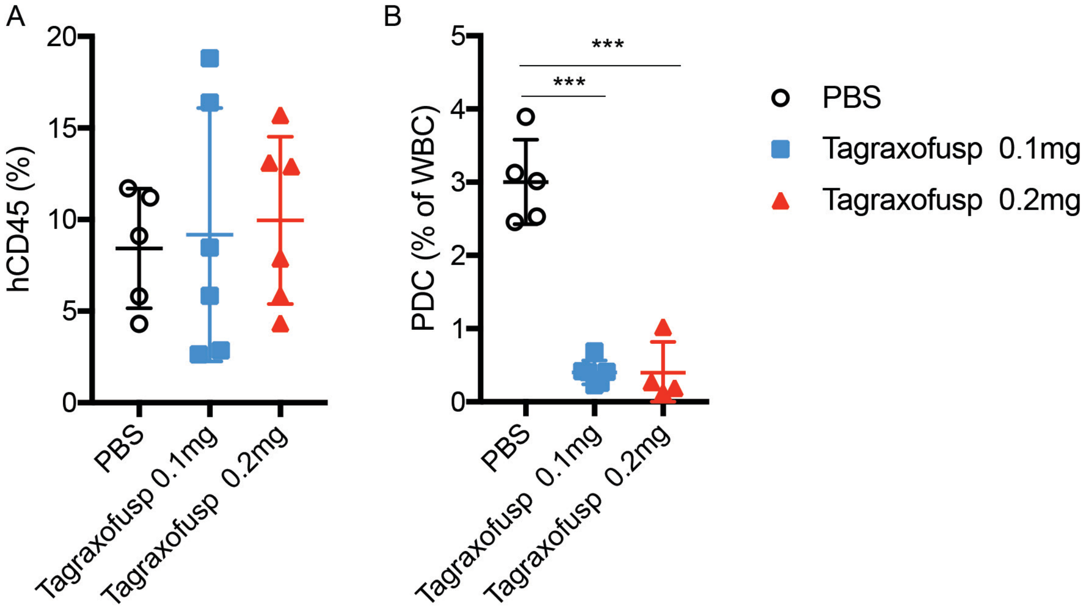


Supplemental Figure 6. Tagraxofusp treatment eliminates pDCs in secondary NSG recipients. A, hCD45 positive cells in bone marrow prior to treatment were comparable between 3 cohorts. B, pDC proportions in total WBC of peripheral blood were nearly eliminated by one cycle of tagraxofusp treatment. mean±SD. ***p<0.0001

Supplemental Table 1 List of antibodies for pDC-AML immunophenotype by flow cytometry

| M1 Tube | M2 Tube | M3 Tube | Intra Tube | pDC1 Tube | pDC2 Tube | pDC3 Tube |
|---|---|---|---|---|---|---|
| CD15 FITC (80H5, BC) | CD64 FITC (22, BC) | CD7 BB515 (M-T701, BD) | cytoplasmic MPO FITC (CLB-MPO-1, BC) | CD303 (BCDA-2) PE (AC144, Miltenyi) | cytoplasmic TdT control FITC (Supertechs) | cytoplasmic TdT FITC (Supertechs) |
| CD33 PE (D3HL60.251, BC) | CD123 PE (9F5, BD) | CD56 PE (N901 [HLDA6], BC) | cytoplasmic CD79a PE (HM47, BC) | CD123 APC (9F5, BD) | CD34 PERCP-CY5.5 (8G12, BD) | CD34 PERCP-CY5.5 (8G12, BD) |
| CD117 PC5 (104D2D1, BC) | CD14 PC5 (RMO052, BC) | CD5 Percp-cy 5.5 (L17F12, BD) | cytoplasmic CD3 APC (UCHT1, BC) | CD56 PC7 (N901 [HLDA6], BC) | CD123 APC (9F5, BD) | CD123 APC (9F5, BD) |
| CD13 PE-CY7 (L138, BD) | CD13 PE-CY7 (L138, BD) | CD34 APC (581, BC) | CD34 PERCP-CY5.5 (8G12, BD) | CD4 PC5.5 (13B8.2, BC) | CD45 APC-H7 (2D1, BD) | CD45 APC-H7 (2D1, BD) |
| CD34 APC (581, BC) | CD34 APC (581, BC) | CD33 PC7 (D3HL60.251, BC) | CD19 PC7 (J3-119, BC) | CD45 APC-H7 (2D1, BD) | HLADR PACIFIC BLUE (Immu-357, BC) | HLADR PACIFIC BLUE (Immu-357, BC) |
| CD71 APC-A700 (YDJ1.2.2, BD) | CD16 APC -A700 (3G8, BC) | CD4 APC-A700 (13B8.2, BC) | CD45 APC-H7 (2D1, BD) | HLADR PACIFIC BLUE (Immu-357, BC) | | |
| CD38 APC-A750 (LS198-4-3, BC) | CD38 APC-A750 (LS198-4-3, BC) | CD38 APC-A750 (LS198-4-3, BC) | CD3 BV421 (UCHT1, BD) | | | |
| HLADR PACIFIC BLUE (Immu-357, BC) | HLADR PACIFIC BLUE (Immu-357, BC) | CD2 BV421 (RPA-2.10, BD) | | | | |
| CD45 V500c (2D1, BD) | CD45 V500c (2D1, BD) | CD45 V500c (2D1, BD) | | | | |
| CD19 BV605 (HIB19, BioL) | CD11b BV605 (1CRF44, BioL) | CD25 BV605 (BC96, BioL) | | | | |

BC: Beckman Coulter, Miami, FL; BD: BD Biosciences, San Jose, CA; BioL: BioLegend, San Jose, CA; Miltenyi: Miltenyi Biotec, San Jose, CA; Supertechs: Supertechs, Inc., Rockville, MD

Supplemental Table 2 Immunophenotype of pDC in pDC-AML

| Markers | pDC-AML at diagnosis (n = 26) (FC available n = 25) | pDC-AML late (n=16) (FC available n = 16) | pDC-AML (combined) (n=42) (FC available n = 41) | BPDCN[a] | Normal pDCs[a] |
|---|---|---|---|---|---|
| CD4 | 20/20 (100%) | 9/9 (100%) | 29/29 (100%) | 37/37 (100%) | 21/21 (100%) |
| CD123 | 25/25 (100%) | 15/15 (100%) | 41/41 (100%) | 37/37 (100%) | 21/21 (100%) |
| HLA-DR | 25/25 (100%) | 15/15 (100%) | 41/41 (100%) | 37/37 (100%) | 21/21 (100%) |
| CD303 | 5/8 (63%) | 2/2 (100%) | 7/10 (70%) | 7/16 (44%) | 21/21 (100%) |
| CD56 | 4/20 (20%) | 1/9 (11%) | 5/29 (17%) | 36/37 (97%) | Small population |
| TCL-1 | 1/7 (14%) | 2/2 (100%) | 3/9 (33%) | | |
| TdT | 4/8 (50%) | 0/2 0%) | 4/10 (40%) | 4/16 (25%) | |
| CD34 | 14/25 (56%) | 11/16 (69%) | 25/41 (61%) | 0/27 (0) | |
| CD38 | 24/25 (96%) | 16/16 (100%) | 40/41 (98%) | 30/34 (88%) | 21/21 (100%) |
| CD2 | 6/20 (30%) | 2/9 (22%) | 8/29 (28%) | 5/27 (19%) | 21/21 (100%) |
| CD5 | 3/20 (15%) | 2/9 (22%) | 5/29 (17%) | 1/30 (3%) | |
| CD7 | 8/20 (40%) | 4/9 (44%) | 12/29 (41%) | 21/33 (64%) | 21/21 (100%) |
| CD13 | 9/25 (36%) | 8/16 (50%) | 17/41 (41%) | 0/30 (0) | |
| CD14 | 0/25 (0%) | 0/16 (0%) | 0/41 (0%) | | |
| CD33 | 4/21 (19%) | 7/12 (58%) | 11/33 (33%) | 16/33 (48%) | 21/21 (100%) |
| CD64 | 0/25 (0%) | 1/16 (6%) | 1/41 (2%) | | |

[a] Data were extracted from Wang, W., et al. (2020). "Immunophenotypic characterization of reactive and neoplastic plasmacytoid dendritic cells permits establishment of a 10-color flow cytometric panel for initial workup and residual disease evaluation of blastic plasmacytoid dendritic cell neoplasm." Haematologica: haematol.2020.247569.

Supplemental Table 3 Immunophenotype of blasts in pDC-AML

| **Markers** | **pDC-AML at diagnosis (n = 26) (FC available n = 25)** | **pDC-AML late (n=16) (FC available n = 16)** | **pDC-AML (combined) (n=42) (FC available n = 41)** |
|---|---|---|---|
| Cross lineage antigen expression | 6 (24%) | 7 (44%) | 13 (32%) |
| T-cell markers[a] (≥2 aberrantly expressed) | 4 (16%) | 5 (31%) | 9 (22%) |
| B-cell markers[b] (≥2 aberrantly expressed) | 2 (8%) | 2 (13%) | 4 (10%) |
| Meets MPAL criteria | 5 (20%) | 3 (19%) | 8 (20%) |
| T-cell markers<br>Number abnormal / number tested (%) | | | |
| CD2 | 4/25 (16%) | 2/16 (13%) | 6/41 (15%) |
| CD3 (cytoplasmic) | 0/19 (0%) | 2/5 (40%) | 2/24 (8%) |
| CD5 | 2/25 (8%) | 2/16 (13%) | 4/41 (10%) |
| CD7 | 10/25 (40%) | 9/16 (56%) | 19/41 (46%) |
| B-cell markers | | | |
| CD10 | 0/24 (0%) | 2/5 (40%) | 2/29 (7%) |
| CD19 | 7/25 (28%) | 4/16 (25%) | 11/41 (27%) |
| CD79a (cytoplasmic) | 4/14 (29%) | 2/5 (40%) | 6/19 (32%) |
| Other markers | | | |
| CD4 | 2/25 (8%) | 0/16 (0%) | 2/41 (5%) |
| CD11b | 11/25 (44%) | 4/16 (25%) | 15/41 (37%) |
| CD13 | 24/25 (96%) | 11/16 (69%) | 35/41 (85%) |
| CD14 | 0/25 (0%) | 0/16 (0%) | 0/41 (0%) |
| CD15 | 13/25 (52%) | 2/16 (13%) | 15/41 (37%) |
| CD25 | 10/25 (40%) | 5/16 (31%) | 15/41 (37%) |
| CD33 | 23/25 (92%) | 14/16 (88%) | 37/41 (90%) |
| CD34 | 17/25 (68%) | 12/16 (75%) | 19/41 (46%) |
| CD38 | 14/25 (56%) | 8/16 (50%) | 22/41 (54%) |
| CD56 | 3/25 (12%) | 3/16 (19%) | 6/41 (15%) |
| CD64 | 4/25 (16%) | 2/16 (13%) | 6/41 (15%) |
| CD117 | 24/25 (96%) | 14/16 (88%) | 38/41 (93%) |
| CD123 | 21/25 (84%) | 10/16 (63%) | 31/41 (76%) |
| HLA-DR | 20/25 (80%) | 12/16 (75%) | 32/41 (78%) |

Supplemental Table 4 Clinicopathologic features of 42 patients with pDC-AML

| Patient | Sex | Age | Diagnosis (WHO Category) | Known MDS | Known CMML | Extramedullary Disease | Flowp pDCs (%) | Karyotype | Survival status | Overall Survival |
|---|---|---|---|---|---|---|---|---|---|---|
| Patient 01 | M | 78 | AML-MRC | yes | no | No | 2.3% | 45,XY,-7[4]/46,XY[17] | Died | 15.9 |
| Patient 02 | M | 77 | AML NOS | no | no | No | 3.5% | 47,XY,+13[2]/48,idem,+19[14]/46,XY[4] | Died | 13.4 |
| Patient 03 | F | 73 | AML NOS | no | no | Leukemia cutis | 3.7% | not performed | Died | 6.7 |
| Patient 04 | M | 67 | AML NOS | no | no | Leukemia cutis | 14.0% | 47,XY,+13 [14]/46,XY [6] | Alive | 41.7 |
| Patient 05 | M | 80 | AML NOS | yes | no | No | 3.7% | 45,X,-Y[19]/46,XY[1] | Died | 24.3 |
| Patient 06 | M | 72 | AML NOS | no | no | No | 20.5% | 46,XY[20] | Died | 1.0 |
| Patient 07 | M | 77 | AML NOS | no | no | No | 5.1% | 45,X,-Y,del(2)(p13p21)[21] | Died | 3.5 |
| Patient 08 | M | 47 | AML-MRC | no | no | Leukemia cutis | 5.5% | 46,XY,add(7)(q32)[6]/ 92,XXYY[5]/46,XY[9] | Alive | 9.8 |
| Patient 09 | M | 74 | AML-MRC | no | no | No | 35.9% | 46,XY,del(20)(q11.2q13.1)[16]/46,idem,der(7)t(1;7)(q21;q22)[5] | Died | 18.8 |
| Patient 10 | M | 72 | AML-MRC | no | no | No | 5.8% | 48,XY,+8,+21[3]/46,XY[17] | Died | 2.8 |
| Patient 11 | M | 79 | AML-MRC | yes | no | No | 14.0% | 46,XY[20] | Alive | 6.9 |
| Patient 12 | F | 62 | AML-MRC | yes | no | No | 4.1% | 46,XX,t(8;21)(q11.2;q22)[2]/46,idem,del(3)(q12q29),del(17)(p11.2)[4]/46,XX[13] | Alive | 0.0 |
| Patient 13 | F | 44 | AML NOS | no | no | No | 7.7% | 46,XX[20] | Alive | 10.6 |
| Patient 14 | M | 72 | t-AML | no | no | No | 2.6% | 46,XY[1]/Outside 11/12/2018: 45,X-Y[3]/[46,XY[17] | Alive | 5.3 |
| Patient 15 | F | 76 | AML-MRC | yes | no | No | 9.8% | 46,XX,t(15;21)(q21;q22)[17]/46,XX[3] | Died | 4.9 |
| Patient 16 | M | 74 | AML NOS | no | no | No | 2.5% | 46,XY[20] | Died | 36.3 |
| Patient 17 | F | 14 | t-AML | tMDS | no | No | 10.4% | 46,XX,del(5)(q22q35) [20] | Died | 7.1 |
| Patient 18 | M | 61 | AML NOS | no | no | No | 12.9% | 46,XY[20] | Alive | 8.7 |
| Patient 19 | M | 68 | AML NOS | no | no | No | 8.1% | 79-88<4n>,XXYY,-3,del(5)(q23q35)x2,-7,-7,-11,-16,-17,-21,inc[cp2]/46,XY[12] | Died | 7.4 |
| Patient 20 | M | 75 | AML-MRC | yes | no | Leukemia cutis | 8.2% | 46,XY[20] | Died | 9.8 |
| Patient 21 | M | 81 | AML NOS | no | no | No | 5.4% | 47,XY,t(4;6)(q31;q27),+13[1]/90~92,idemx2,inc[2]/46,XY[11] | Died | 1.7 |
| Patient 22 | M | 70 | AML-MRC | yes | no | No | 10.0% | 46,XY[20] | Died | 42.6 |
| Patient 23 | M | 65 | AML-MRC | yes | no | No | 4.8% | 46,XY,inv(4)(p15q21) [4]/46,XY [5] | Died | 22.1 |
| Patient 24 | F | 66 | AML NOS | yes | yes | Lymph Node | 13.2% | 47,XX,+8[12]/46,XX[8] | Died | 10.2 |
| Patient 25 | F | 77 | AML-MRC | no | no | No | 2.1% | 46,XX[20] | Alive | 17.9 |
| Patient 26 | M | 63 | AML-MRC | yes | yes | Leukemia cutis | | 46,XY[20] | Alive | 11.0 |
| Patient 27 | M | 71 | AML NOS | no | no | No | 6.3% | 46,XY[20] | Died | 21.0 |
| Patient 28 | M | 58 | AML NOS | no | no | No | 26.3% | 46,XY[15] | Died | 3.8 |
| Patient 29 | M | 56 | AML MRC | no | no | Lymph Node | 3.5% | 46,XY,+X,-7[16]/46,XY[4] | Alive | 5.6 |
| Patient 30 | M | 65 | AML MRC | yes | no | No | 5.9% | 46,XY,del(9)(q22q32)[6]/46,idem,t(18;19)(q21;p13.3)[1]/46,idem,del(1)(q32)[1]/46,XY,del(9)(q12q34)[1]/46,XY[11] | Alive | 16.6 |
| Patient 31 | F | 68 | AML NOS | no | no | No | 4.6% | karyotype failure | Died | 30.8 |
| Patient 32 | M | 54 | t-AML | no | no | No | 3.0% | 45,XY,-7[14] | Died | 14.1 |
| Patient 33 | M | 15 | AML MRC | yes | no | No | 7.7% | 46,XY,dup(1)(q22q42)[8]/46,idem,add(6)(p23)[6] | Died | 46.3 |
| Patient 34 | M | 76 | AML MRC | yes | no | No | 2.2% | 46,XY[14] use this was 46,XY,+1,der(1;7)(q10;p10)[8/12] | Died | 51.8 |
| Patient 35 | M | 78 | AML MRC | yes | no | No | 2.5% | 46,XY[20] | Died | 7.3 |
| Patient 36 | M | 9 | AML MRC | no | no | No | 2.9% | 45,XY,-7 [20] | Died | 96.6 |
| Patient 37 | M | 59 | t-AML | no | no | No | 3.3% | 46,XY[20] | Alive | 22.6 |

| | | | | | | | | | | |
|---|---|---|---|---|---|---|---|---|---|---|
| Patient 38 | M | 45 | AML NOS | no | no | No | 2.2% | 46,XY[20] | Alive | 43.6 |
| Patient 39 | F | 76 | AML NOS | no | no | No | 9.2% | 46,XX[20] | Died | 13.0 |
| Patient 40 | F | 65 | AML NOS | no | no | No | 2.5% | 46,XX[20] | Died | 14.6 |
| Patient 41 | M | 29 | AML NOS | no | no | No | 4.4% | 46,XY,?del(12)(p12)[3]/46,XY[22] | Alive | 51.0 |
| Patient 42 | F | 31 | AML w NPM1 | no | no | No | 2.8% | 46,XY[20] | Alive | 13.3 |

Patients 1-26 were pDC-AML at diagnosis. Patients 27-42 were pDC-AML late (after diagnosis).

Supplemental Table 5. Mutational annotation of pDC-AML

| PT# | Ploatform | Tm | chr | start pos | Gene | TxID | cDNA_annotation | AA_annotation | VAF | Annotation |
|---|---|---|---|---|---|---|---|---|---|---|
| **1** | RD | D | 20 | 31022615 | ASXL1 | NM_015338 | c.2101delC | p.P701fs | 18.80% | Likely ONCOGENIC |
| **1** | RD | D | X | 133551212 | PHF6 | NM_032458 | c.848G>T | p.C283F | 31.70% | Likely ONCOGENIC |
| **1** | RD | D | 21 | 36259172 | RUNX1 | NM_001754 | c.319C>T | p.R107C | 14.40% | Likely ONCOGENIC |
| **1** | RD | D | X | 133551305 | PHF6 | NM_032458 | c.941T>C | p.I314T | 15.00% | Likely ONCOGENIC |
| **1** | RD | D | 4 | 106157989 | TET2 | NM_001127208 | c.2890C>T | p.Q964X | 18.40% | Likely ONCOGENIC |
| **1** | RD | F#1 | 20 | 31022615 | ASXL1 | NM_015338 | c.2101delC | p.P701fs | 41.50% | Likely ONCOGENIC |
| **1** | RD | F#1 | X | 133551212 | PHF6 | NM_032458 | c.848G>T | p.C283F | 70.40% | Likely ONCOGENIC |
| **1** | RD | F#1 | 21 | 36259172 | RUNX1 | NM_001754 | c.319C>T | p.R107C | 32.90% | Likely ONCOGENIC |
| **1** | RD | F#1 | 12 | 111856098 | SH2B3 | NM_005475 | c.149G>A | p.R50Q | 6.10% | UNKNOWN |
| **1** | RD | F#1 | 4 | 106157989 | TET2 | NM_001127208 | c.2890C>T | p.Q964X | 32.80% | Likely ONCOGENIC |
| **2** | RD | D | 2 | 209113112 | IDH1 | NM_005896 | c.395G>A | p.R132H | 13.80% | ONCOGENIC |
| **2** | RD | D | 21 | 36171683 | RUNX1 | NM_001754 | c.881delC | p.P294fs | 75.20% | Likely ONCOGENIC |
| **3** | RD | D | 21 | 36231864 | RUNX1 | NM_001754 | c.520A>C | p.T174P | 36.90% | Likely ONCOGENIC |
| **4** | IMPACT | D | 21 | 36164571 | RUNX1 | NM_001754 | c.1303dupG | p.A435Gfs*165 | 6.50% | Likely ONCOGENIC |
| **5** | RD | D | 20 | 31022989 | ASXL1 | NM_015338 | c.2475_2476delAG | p.G826Nfs*6 | 24.73% | Likely ONCOGENIC |
| **6** | IMPACT | D | 21 | 36259198 | RUNX1 | NM_001754 | c.292delC | p.L98Sfs*24 | 53.80% | Likely ONCOGENIC |
| **6** | IMPACT | D | 2 | 25467449 | DNMT3A | NM_022552 | c.1627G>T | p.G543C | 42.70% | Likely ONCOGENIC |
| **6** | IMPACT | D | 2 | 25463539 | DNMT3A | NM_022552 | c.2142_2143insGGTCAAACAAGCCC | p.I715Gfs*69 | 19.70% | Likely ONCOGENIC |
| **7** | RD | D | 21 | 36259138 | RUNX1 | NM_001754 | c.351+2T>C | Splicing | 42.62% | Likely ONCOGENIC |
| **7** | RD | D | 21 | 36259186 | RUNX1 | NM_001754 | c.305T>C | p.L102P | 10.31% | Likely ONCOGENIC |
| **7** | RD | D | 17 | 30274671 | SUZ12 | NM_015355 | c.422T>C | p.V141A | 55.05% | UNKNOWN |
| **8** | IMPACT | D | 1 | 115258747 | NRAS | NM_002524 | c.35G>C | p.G12A | 2.40% | ONCOGENIC |
| **8** | IMPACT | D | 21 | 36164685 | RUNX1 | NM_001754 | c.1189delC | p.Q397Kfs*197 | 22.70% | Likely ONCOGENIC |
| **8** | IMPACT | D | 17 | 74732959 | SRSF2 | NM_003016 | c.284C>T | p.P95L | 6.20% | ONCOGENIC |
| **8** | IMPACT | D | 12 | 112888199 | PTPN11 | NM_002834 | c.215C>G | p.A72G | 16.70% | Likely ONCOGENIC |
| **9** | IMPACT | D | 2 | 25966453 | ASXL2 | NM_015338 | c.2752_2753delinsGGT | p.S918Gfs*8 | 17.00% | Likely ONCOGENIC |
| **9** | IMPACT | D | X | 133511704 | PHF6 | NM_032458 | c.59_70delinsTT | p.C20Ffs*10 | 33.40% | Likely ONCOGENIC |
| **10** | RD | D | 2 | 198267371 | SF3B1 | NM_012433 | c.1986C>A | p.H662Q | 16.10% | ONCOGENIC |

| | | | | | | | | | | |
|---|---|---|---|---|---|---|---|---|---|---|
| **10** | RD | D | 7 | 148514354 | EZH2 | NM_004456 | c.1370G>A | p.C457Y | 38.30% | UNKNOWN |
| **10** | RD | D | 20 | 31023408 | ASXL1 | NM_015338 | c.2893C>T | p.R965* | 18.90% | ONCOGENIC |
| **10** | RD | D | 21 | 36231783 | RUNX1 | NM_001754 | c.601C>T | p.R201* | 18.30% | ONCOGENIC |
| **10** | RD | F#1 | 2 | 198267371 | SF3B1 | NM_012433 | c.1986C>A | p.H662Q | 10.95% | ONCOGENIC |
| **10** | RD | F#1 | 7 | 148514354 | EZH2 | NM_004456 | c.1370G>A | p.C457Y | 7.60% | UNKNOWN |
| **10** | RD | F#1 | 20 | 31023408 | ASXL1 | NM_015338 | c.2893C>T | p.R965* | 1.00% | ONCOGENIC |
| **10** | RD | F#1 | 9 | 5073770 | JAK2 | NM_004972 | c.G1849T | p.V617F | 0.10% | ONCOGENIC |
| **10** | RD | F#2 | 2 | 198267371 | SF3B1 | NM_012433 | c.1986C>A | p.H662Q | 25.37% | ONCOGENIC |
| **10** | RD | F#2 | 7 | 148514354 | EZH2 | NM_004456 | c.1370G>A | p.C457Y | 44.25% | UNKNOWN |
| **10** | RD | F#2 | 20 | 31023408 | ASXL1 | NM_015338 | c.2893C>T | p.R965* | 18.04% | ONCOGENIC |
| **10** | RD | F#2 | 21 | 36231783 | RUNX1 | NM_001754 | c.601C>T | p.R201* | 7.67% | ONCOGENIC |
| **10** | RD | F#3 | 2 | 198267371 | SF3B1 | NM_012433 | c.1986C>A | p.H662Q | 38.70% | ONCOGENIC |
| **10** | RD | F#3 | 7 | 148514354 | EZH2 | NM_004456 | c.1370G>A | p.C457Y | 70.51% | UNKNOWN |
| **10** | RD | F#3 | 20 | 31023408 | ASXL1 | NM_015338 | c.2893C>T | p.R965* | 26.94% | ONCOGENIC |
| **10** | RD | F#3 | 21 | 36231783 | RUNX1 | NM_001754 | c.601C>T | p.R201* | 37.43% | ONCOGENIC |
| **10** | RD | F#3 | 21 | 36259152 | RUNX1 | NM_001754 | c.338_339insGCAG | p.I114Qfs*25 | 15.35% | Likely ONCOGENIC |
| **10** | RD | F#4 | 2 | 198267371 | SF3B1 | NM_012433 | c.1986C>A | p.H662Q | 50.15% | ONCOGENIC |
| **10** | RD | F#4 | 7 | 148514354 | EZH2 | NM_004456 | c.1370G>A | p.C457Y | 92.35% | UNKNOWN |
| **10** | RD | F#4 | 13 | 28608260 | FLT3 | NM_004119 | c.1795_1796insCCCTTGATTTCAGAGAATATGAAT | p.E598_Y599insSLDFREYE | 3.10% | ONCOGENIC |
| **10** | RD | F#4 | 20 | 31023408 | ASXL1 | NM_015338 | c.2893C>T | p.R965* | 62.16% | ONCOGENIC |
| **10** | RD | F#4 | 21 | 36231783 | RUNX1 | NM_001754 | c.601C>T | p.R201* | 68.92% | ONCOGENIC |
| **10** | RD | F#4 | 21 | 36259152 | RUNX1 | NM_001754 | c.338_339insGCAG | p.I114Qfs*25 | 48.17% | Likely ONCOGENIC |
| **11** | RD | D | 7 | 148544353 | EZH2 | NM_004456 | c.26_36delinsG | p.E9Gfs*8 | 24.79% | UNKNOWN |
| **11** | RD | D | 7 | 148508721 | EZH2 | NM_004456 | c.1943G>C | p.G648A | 6.56% | UNKNOWN |
| **11** | RD | D | 9 | 139399237 | NOTCH1 | NM_017617 | c.4906G>A | p.E1636K | 66.46% | UNKNOWN |
| **11** | RD | D | 20 | 31022497 | ASXL1 | NM_015338 | c.1982_1990delinsCTCT | p.R661Tfs*5 | 25.63% | Likely ONCOGENIC |
| **11** | RD | D | X | 123211892 | STAG2 | NM_001042749 | c.2759_2760insAA | p.L921Ifs*11 | 26.49% | Likely ONCOGENIC |
| **11** | RD | F#1 | 7 | 148544353 | EZH2 | NM_004456 | c.26_36delinsG | p.E9Gfs*8 | 42.16% | UNKNOWN |
| **11** | RD | F#1 | 7 | 14850872 | EZH2 | NM_004456 | c.1943G>C | p.G648A | 31.14% | UNKNOWN |

| | | | | | | | | | | |
|---|---|---|---|---|---|---|---|---|---|---|
| | | | | 1 | | | | | | |
| **11** | RD | F#1 | 9 | 139399237 | NOTCH1 | NM_017617 | c.4906G>A | p.E1636K | 66.01% | UNKNOWN |
| **11** | RD | F#1 | 20 | 31022497 | ASXL1 | NM_015338 | c.1982_1990delinsCTCT | p.R661Tfs*5 | 41.13% | Likely ONCOGENIC |
| **11** | RD | F#1 | X | 123211892 | STAG2 | NM_001042749 | c.2759_2760insAA | p.L921Ifs*11 | 41.95% | Likely ONCOGENIC |
| **12** | IMPACT | D | 21 | 36171718 | RUNX1 | NM_001754 | c.847C>T | p.Q283* | 76.00% | Likely ONCOGENIC |
| **12** | IMPACT | D | 13 | 28592642 | FLT3 | NM_004119 | c.2503G>T | p.D835Y | 54.10% | ONCOGENIC |
| **12** | IMPACT | D | 4 | 106158000 | TET2 | NM_001127208 | c.2905delC | p.Q969Kfs*38 | 42.90% | Likely ONCOGENIC |
| **12** | IMPACT | D | 17 | 74732959 | SRSF2 | NM_003016 | c.284C>A | p.P95H | 46.80% | ONCOGENIC |
| **12** | IMPACT | D | 17 | 7577099 | TP53 | NM_000546 | c.839G>C | p.R280T | 3.50% | ONCOGENIC |
| **12** | IMPACT | D | 1 | 115256529 | NRAS | NM_002524 | c.182A>G | p.Q61R | 38.50% | ONCOGENIC |
| **12** | IMPACT | D | X | 39932146 | BCOR | NM_001123385 | c.2452_2453insGTCTCCTG | p.D818Gfs*41 | 70.80% | Likely ONCOGENIC |
| **12** | IMPACT | D | 18 | 42531913 | SETBP1 | NM_015559 | c.2608G>A | p.G870S | 4.70% | ONCOGENIC |
| **12** | IMPACT | D | 2 | 198267371 | SF3B1 | NM_012433 | c.1986C>A | p.H662Q | 47.70% | Likely ONCOGENIC |
| **12** | IMPACT | D | 20 | 31022468 | ASXL1 | NM_015338 | c.1955dupG | p.G653Rfs*5 | 40.00% | Likely ONCOGENIC |
| **13** | RD | D | 17 | 74732935 | SRSF2 | NM_003016 | c.284_307delCCCCGGACTCACACCACAGCCGCC | p.P95_R102del | 39.48% | ONCOGENIC |
| **13** | RD | D | X | 39922106 | BCOR | NM_001123385 | c.4066A>T | p.K1356* | 65.99% | ONCOGENIC |
| **13** | RD | D | 21 | 36252878 | RUNX1 | NM_001754 | c.484A>G | p.R162G | 58.40% | Likely ONCOGENIC |
| **14** | RD | D | 1 | 115258747 | NRAS | NM_002524 | c.35G>A | p.G12D | 36.40% | ONCOGENIC |
| **14** | RD | D | 21 | 36231791 | RUNX1 | NM_001754 | c.593A>G | p.D198G | 44.40% | Likely ONCOGENIC |
| **15** | RD | D | 2 | 25464525 | DNMT3A | NM_022552 | c.1988C>T | p.S663L | 11.00% | UNKNOWN |
| **15** | RD | D | 21 | 44524456 | U2AF1 | NM_006758 | c.101C>T | p.S34F | 37.00% | ONCOGENIC |
| **15** | RD | D | 21 | 36259175 | RUNX1 | NM_001754 | c.314_315delAC | p.H105Lfs*32 | 26.50% | Likely ONCOGENIC |
| **16** | IMPACT | D | X | 133551305 | PHF6 | NM_032458 | c.941T>C | p.I314T | 26.19% | Likely ONCOGENIC |
| **16** | IMPACT | D | 13 | 28608320 | FLT3 | NM_004119 | c.1736T>A | p.V579E | 18.84% | Likely ONCOGENIC |
| **16** | IMPACT | D | 21 | 36252994 | RUNX1 | NM_001754 | c.366_367dupGG | p.D123Gfs*11 | 44.85% | Likely ONCOGENIC |
| **16** | IMPACT | D | X | 133551305 | PHF6 | NM_032458 | c.941T>C | p.I314T | 63.88% | Likely ONCOGENIC |
| **17** | RD | D | 20 | 31022547 | ASXL1 | NM_015338 | c.2036delG | p.G679Efs*24 | 12.68% | Likely ONCOGENIC |
| **17** | RD | D | 21 | 36259198 | RUNX1 | NM_001754 | c.292delC | p.L98Sfs*24 | 5.41% | Likely ONCOGENIC |
| **17** | RD | F#1 | 20 | 31022547 | ASXL1 | NM_015338 | c.2036delG | p.G679Efs*24 | 28.29% | Likely ONCOGENIC |

| | | | | | | | | | | |
|---|---|---|---|---|---|---|---|---|---|---|
| **17** | RD | F#1 | 21 | 36259198 | RUNX1 | NM_001754 | c.292delC | p.L98Sfs*24 | 25.53% | Likely ONCOGENIC |
| **17** | RD | F#1 | 21 | 36252865 | RUNX1 | NM_001754 | c.497G>T | p.R166L | 7.11% | LIKELY ONCOGENIC |
| **17** | RD | F#1 | 21 | 36252877 | RUNX1 | NM_001754 | c.485G>A | p.R162K | 10.48% | Likely ONCOGENIC |
| **17** | RD | F#1 | X | 44938515 | KDM6A | NM_021140 | c.3063G>A | p.W1021* | 31.87% | UNKNOWN |
| **17** | RD | F#1 | 4 | 106156042 | TET2 | NM_001127208 | c.945delC | p.Q317Rfs*30 | 11.50% | Likely ONCOGENIC |
| **17** | RD | F#1 | 4 | 106164866 | TET2 | NM_001127208 | c.3734A>G | p.Y1245C | 12.25% | Likely ONCOGENIC |
| **17** | RD | F#1 | 21 | 36206762 | RUNX1 | NM_001754 | c.749_750insGTTGACCC | p.A251Lfs*6 | 58.92% | Likely ONCOGENIC |
| **17** | RD | F#1 | X | 39931909 | BCOR | NM_001123385 | c.2690C>A | p.S897* | 69.72% | ONCOGENIC |
| **18** | IMPACT | D | 1 | 115258674 | NRAS | NM_002524 | c.108A>G | p.I36M | 46.60% | UNKNOWN |
| **18** | IMPACT | D | 20 | 31022441 | ASXL1 | NM_015338 | c.1934dupG | p.G646Wfs*12 | 35.10% | ONCOGENIC |
| **18** | IMPACT | D | 21 | 36171751 | RUNX1 | NM_001754 | c.814C>T | p.Q272* | 34.90% | Likely ONCOGENIC |
| **18** | IMPACT | D | X | 39922119 | BCOR | NM_001123385 | c.4051_4052delAC | p.T1351Rfs*57 | 86.40% | Likely ONCOGENIC |
| **18** | IMPACT | D | X | 133559265 | PHF6 | NM_032458 | c.1003A>T | p.R335* | 91.60% | ONCOGENIC |
| **18** | IMPACT | D | X | 39922124 | BCOR | NM_001123385 | c.4048T>A | p.Y1350N | 87.90% | UNKNOWN |
| **19** | IMPACT | D | 12 | 49428194 | KMT2D | NM_003482 | c.10505_10506insTTTACCC | p.N3503Lfs*4 | 7.62% | Likely ONCOGENIC |
| **19** | IMPACT | D | 2 | 198267705 | SF3B1 | NM_012433 | c.1774G>A | p.E592K | 18.93% | UNKNOWN |
| **19** | IMPACT | D | 20 | 31022402 | ASXL1 | NM_015338 | c.1900_1922delAGAGAGGCGGCCACCACTGCCAT | p.E635Rfs*15 | 19.79% | Likely ONCOGENIC |
| **19** | IMPACT | D | 21 | 36252865 | RUNX1 | NM_001754 | c.497G>A | p.R166Q | 17.55% | LIKELY ONCOGENIC |
| **19** | IMPACT | D | 4 | 106190860 | TET2 | NM_001127208 | c.4138C>T | p.H1380Y | 17.10% | Likely ONCOGENIC |
| **19** | IMPACT | D | 8 | 117874082 | RAD21 | NM_006265 | c.371_372insTGAGGGCGGA | p.L124Ffs*6 | 5.66% | UNKNOWN |
| **19** | IMPACT | D | X | 133547940 | PHF6 | NM_032458 | c.673C>T | p.R225* | 20.24% | ONCOGENIC |
| **20** | IMPACT | D | 1 | 115258747 | NRAS | NM_002524 | c.35G>T | p.G12V | 13.40% | ONCOGENIC |
| **20** | IMPACT | D | 12 | 25398285 | KRAS | NM_033360 | c.34G>C | p.G12R | 2.40% | ONCOGENIC |
| **20** | IMPACT | D | 4 | 106193931 | TET2 | NM_001127208 | c.4393C>T | p.R1465* | 16.00% | Likely ONCOGENIC |
| **20** | IMPACT | D | 4 | 106157326 | TET2 | NM_001127208 | c.2227C>T | p.Q743* | 52.90% | Likely ONCOGENIC |
| **20** | IMPACT | D | 11 | 119148919 | CBL | NM_005188 | c.1139T>C | p.L380P | 21.50% | Likely ONCOGENIC |
| **20** | IMPACT | D | 17 | 74732959 | SRSF2 | NM_003016 | c.284C>T | p.P95L | 34.00% | Likely ONCOGENIC |
| **21** | OSH | D | 1 | 115258747 | NRAS | NM_002524 | c.35G>T | p.G12V | 37.00% | ONCOGENIC |

| | | | | | | | | | | |
|---|---|---|---|---|---|---|---|---|---|---|
| **21** | OSH | D | 21 | 36171607 | RUNX1 | NM_001754 | c.958C>T | p.R320* | 17.00% | Likely ONCOGENIC |
| **21** | OSH | D | 21 | 36259172 | RUNX1 | NM_001754 | c.319C>T | p.R107C | 16.50% | Likely ONCOGENIC |
| **21** | OSH | D | 17 | 74732959 | SRSF2 | NM_003016 | c.284C>A | p.P95H | 42.00% | ONCOGENIC |
| **21** | OSH | D | 20 | 31022402 | ASXL1 | NM_015338 | c.1900_1922delAGAGA GGCGGCCACCACTGCC AT | p.E635fs*15 | 26.00% | Likely ONCOGENIC |
| **22** | RD | D | 2 | 19826735 9 | SF3B1 | NM_012433 | c.1998G>T | p.K666N | 46.90% | Likely ONCOGENIC |
| **22** | RD | D | 21 | 36231774 | RUNX1 | NM_001754 | c.610C>T | p.R204* | 50.00% | Likely ONCOGENIC |
| **22** | RD | D | 13 | 28592642 | FLT3 | NM_004119 | c.2503G>T | p.D835Y | 2.10% | ONCOGENIC |
| **22** | RD | D | 2 | 25469964 | DNMT3 A | NM_022552 | c.1078A>G | p.N360D | 10.00% | UNKNOWN |
| **23** | IMPACT | D | 21 | 36231782 | RUNX1 | NM_001754 | c.602G>A | p.R201Q | 14.40% | ONCOGENIC |
| **23** | IMPACT | D | 2 | 25464537 | DNMT3 A | NM_022552 | c.1976G>A | p.R659H | 31.80% | Likely ONCOGENIC |
| **23** | IMPACT | D | 2 | 25468159 | DNMT3 A | NM_022552 | c.1513_1516delGAAC | p.E505Tfs*145 | 19.50% | Likely ONCOGENIC |
| **23** | IMPACT | D | 21 | 44514777 | U2AF1 | NM_006758 | c.470A>G | p.Q157R | 13.60% | ONCOGENIC |
| **23** | IMPACT | D | 14 | 81558933 | TSHR | NM_000369 | c.526T>C | p.C176R | 14.50% | UNKNOWN |
| **23** | IMPACT | D | 17 | 74732959 | SRSF2 | NM_003016 | c.284C>A | p.P95H | 15.30% | ONCOGENIC |
| **24** | RD | D | 2 | 25457243 | DNMT3 A | NM_022552 | c.C2644T | p.R882C | 36.40% | ONCOGENIC |
| **24** | RD | D | 15 | 90631934 | IDH2 | NM_002168 | c.G419A | p.R140Q | 34.80% | ONCOGENIC |
| **24** | RD | D | 21 | 36171710 | RUNX1 | NM_001754 | c.855C>G | p.Y285X | 28.70% | ONCOGENIC |
| **24** | RD | F#1 | 2 | 25457243 | DNMT3 A | NM_022552 | c.C2644T | p.R882C | 23.70% | ONCOGENIC |
| **24** | RD | F#1 | 15 | 90631934 | IDH2 | NM_002168 | c.G419A | p.R140Q | 33.80% | ONCOGENIC |
| **24** | RD | F#1 | 21 | 36171710 | RUNX1 | NM_001754 | c.855C>G | p.Y285X | 16.70% | ONCOGENIC |
| **24** | RD | F#1 | 21 | 36252954 | RUNX1 | NM_001754 | c.408T>G | p.N136K | 33.20% | Likely ONCOGENIC |
| **24** | RD | F#2 | 2 | 25457243 | DNMT3 A | NM_022552 | c.C2644T | p.R882C | 41.80% | ONCOGENIC |
| **24** | RD | F#2 | 15 | 90631934 | IDH2 | NM_002168 | c.G419A | p.R140Q | 43.60% | ONCOGENIC |
| **24** | RD | F#2 | 21 | 36171710 | RUNX1 | NM_001754 | c.C855G | p.Y285X | 25.20% | ONCOGENIC |
| **24** | RD | F#2 | 21 | 36252954 | RUNX1 | NM_001754 | c.408T>G | p.N136K | 44.90% | Likely ONCOGENIC |
| **24** | RD | F#3 | 2 | 25457243 | DNMT3 A | NM_022552 | c.C2644T | p.R882C | 46.80% | ONCOGENIC |
| **24** | RD | F#3 | 15 | 90631934 | IDH2 | NM_002168 | c.G419A | p.R140Q | 47.40% | ONCOGENIC |
| **24** | RD | F#3 | 21 | 36171710 | RUNX1 | NM_001754 | c.855C>G | p.Y285X | 24.50% | ONCOGENIC |
| **24** | RD | F#3 | 21 | 36252954 | RUNX1 | NM_001754 | c.408T>G | p.N136K | 48.10% | Likely ONCOGENIC |

| | | | | | | | | | | |
|---|---|---|---|---|---|---|---|---|---|---|
| **24** | RD | F#4 | 2 | 25457243 | DNMT3 A | NM_022552 | c.C2644T | p.R882C | 17.60% | ONCOGENIC |
| **24** | RD | F#4 | 15 | 90631934 | IDH2 | NM_002168 | c.G419A | p.R140Q | 22.20% | ONCOGENIC |
| **24** | RD | F#4 | 21 | 36171710 | RUNX1 | NM_001754 | c.855C>G | p.Y285X | 7.00% | ONCOGENIC |
| **24** | RD | F#4 | 21 | 36252954 | RUNX1 | NM_001754 | c.408T>G | p.N136K | 13.80% | Likely ONCOGENIC |
| **24** | RD | F#5 | 2 | 25457243 | DNMT3 A | NM_022552 | c.C2644T | p.R882C | 46.01% | ONCOGENIC |
| **24** | RD | F#5 | 15 | 90631934 | IDH2 | NM_002168 | c.G419A | p.R140Q | 47.45% | ONCOGENIC |
| **24** | RD | F#5 | 21 | 36252954 | RUNX1 | NM_001754 | c.408T>G | p.N136K | 41.02% | Likely ONCOGENIC |
| **25** | RD | D | 2 | 209113113 | IDH1 | NM_005896 | c.394C>T | p.R132C | 41.41% | ONCOGENIC |
| **25** | RD | D | 2 | 25457242 | DNMT3 A | NM_022552 | c.2645G>A | p.R882H | 41.72% | Likely ONCOGENIC |
| **25** | RD | D | 2 | 25464516 | DNMT3 A | NM_022552 | c.1997G>A | p.C666Y | 40.78% | UNKNOWN |
| **25** | RD | D | 11 | 32456486 | WT1 | NM_024426 | c.406C>T | p.P136S | 27.75% | UNKNOWN |
| **25** | RD | D | 11 | 32456248 | WT1 | NM_024426 | c.643_644insCG | p.Q215fs | 34.95% | UNKNOWN |
| **25** | RD | D | 13 | 28592642 | FLT3 | NM_004119 | c.2503G>C | p.D835H | 38.39% | ONCOGENIC |
| **25** | RD | D | 19 | 33792937 | CEBPA | NM_004364 | c.383dupC | p.P129fs | 29.58% | Likely ONCOGENIC |
| **25** | RD | D | 21 | 36252994 | RUNX1 | NM_001754 | c.366_367dupGG | p.D123Gfs*11 | 26.06% | Likely ONCOGENIC |
| **25** | RD | F#1 | 2 | 209113113 | IDH1 | NM_005896 | c.394C>T | p.R132C | 18.99% | ONCOGENIC |
| **25** | RD | F#1 | 2 | 25457242 | DNMT3 A | NM_022552 | c.2645G>A | p.R882H | 26.53% | Likely ONCOGENIC |
| **25** | RD | F#1 | 2 | 25464516 | DNMT3 A | NM_022552 | c.1997G>A | p.C666Y | 27.08% | UNKNOWN |
| **25** | RD | F#1 | 11 | 32456486 | WT1 | NM_024426 | c.406C>T | p.P136S | 33.60% | UNKNOWN |
| **25** | RD | F#1 | 11 | 32456248 | WT1 | NM_024426 | c.643_644insCG | p.Q215fs | 12.85% | UNKNOWN |
| **25** | RD | F#1 | 13 | 28592642 | FLT3 | NM_004119 | c.2503G>C | p.D835H | 11.95% | ONCOGENIC |
| **25** | RD | F#1 | 19 | 33792937 | CEBPA | NM_004364 | c.383dupC | p.P129fs | 9.72% | Likely ONCOGENIC |
| **25** | RD | F#1 | 21 | 36252994 | RUNX1 | NM_001754 | c.366_367dupGG | p.D123Gfs*11 | 10.12% | Likely ONCOGENIC |
| **25** | RD | F#2 | 2 | 209113113 | IDH1 | NM_005896 | c.394C>T | p.R132C | 44.85% | ONCOGENIC |
| **25** | RD | F#2 | 2 | 25457242 | DNMT3 A | NM_022552 | c.2645G>A | p.R882H | 37.30% | Likely ONCOGENIC |
| **25** | RD | F#2 | 2 | 25464516 | DNMT3 A | NM_022552 | c.1997G>A | p.C666Y | 51.25% | UNKNOWN |
| **25** | RD | F#2 | 11 | 32456486 | WT1 | NM_024426 | c.406C>T | p.P136S | 46.80% | UNKNOWN |
| **25** | RD | F#2 | 11 | 32456248 | WT1 | NM_024426 | c.643_644insCG | p.Q215fs | 41.55% | UNKNOWN |
| **25** | RD | F#2 | 13 | 28592642 | FLT3 | NM_004119 | c.2503G>C | p.D835H | 37.46% | ONCOGENIC |
| **25** | RD | F#2 | 19 | 33792937 | CEBPA | NM_004364 | c.383dupC | p.P129fs | 46.26% | Likely ONCOGENIC |

| | | | | | | | | | | |
|---|---|---|---|---|---|---|---|---|---|---|
| **25** | RD | F#2 | 21 | 36252994 | RUNX1 | NM_001754 | c.366_367dupGG | p.D123Gfs*11 | 36.74% | Likely ONCOGENIC |
| **25** | RD | F#3 | 2 | 209113113 | IDH1 | NM_005896 | c.394C>T | p.R132C | 43.29% | ONCOGENIC |
| **25** | RD | F#3 | 2 | 25457242 | DNMT3A | NM_022552 | c.2645G>A | p.R882H | 41.15% | Likely ONCOGENIC |
| **25** | RD | F#3 | 2 | 25464516 | DNMT3A | NM_022552 | c.1997G>A | p.C666Y | 40.46% | UNKNOWN |
| **25** | RD | F#3 | 11 | 32456486 | WT1 | NM_024426 | c.406C>T | p.P136S | 34.81% | UNKNOWN |
| **25** | RD | F#3 | 11 | 32456248 | WT1 | NM_024426 | c.643_644insCG | p.Q215fs | 38.68% | UNKNOWN |
| **25** | RD | F#3 | 13 | 28592642 | FLT3 | NM_004119 | c.2503G>C | p.D835H | 37.28% | ONCOGENIC |
| **25** | RD | F#3 | 19 | 33792937 | CEBPA | NM_004364 | c.383dupC | p.P129fs | 27.99% | Likely ONCOGENIC |
| **25** | RD | F#3 | 21 | 36252994 | RUNX1 | NM_001754 | c.366_367dupGG | p.D123Gfs*11 | 29.87% | Likely ONCOGENIC |
| **25** | RD | F#4 | 2 | 209113113 | IDH1 | NM_005896 | c.394C>T | p.R132C | 44.88% | ONCOGENIC |
| **25** | RD | F#4 | 2 | 25457242 | DNMT3A | NM_022552 | c.2645G>A | p.R882H | 52.79% | Likely ONCOGENIC |
| **25** | RD | F#4 | 2 | 25464516 | DNMT3A | NM_022552 | c.1997G>A | p.C666Y | 43.93% | UNKNOWN |
| **25** | RD | F#4 | 11 | 32456486 | WT1 | NM_024426 | c.406C>T | p.P136S | 29.57% | UNKNOWN |
| **25** | RD | F#4 | 11 | 32456248 | WT1 | NM_024426 | c.643_644insCG | p.Q215fs | 47.15% | UNKNOWN |
| **25** | RD | F#4 | 13 | 28592642 | FLT3 | NM_004119 | c.2503G>C | p.D835H | 46.57% | ONCOGENIC |
| **25** | RD | F#4 | 19 | 33792937 | CEBPA | NM_004364 | c.383dupC | p.P129fs | 44.76% | Likely ONCOGENIC |
| **25** | RD | F#4 | 21 | 36252994 | RUNX1 | NM_001754 | c.366_367dupGG | p.D123Gfs*11 | 32.76% | Likely ONCOGENIC |
| **26** | RD | F#1 | 21 | 36252865 | RUNX1 | NM_001754 | c.497G>A | p.R166Q | 41.20% | Likely ONCOGENIC |
| **27** | RD | F#1 | 2 | 209113113 | IDH1 | NM_005896 | c.394C>A | p.R132S | 29.37% | Likely ONCOGENIC |
| **27** | RD | F#1 | 2 | 25467408 | DNMT3A | NM_022552 | c.1667+1G>A | p.X556_splice | 89.10% | ONCOGENIC |
| **27** | RD | F#1 | 17 | 7577091 | TP53 | NM_000546 | c.847C>T | p.R283C | 65.57% | ONCOGENIC |
| **27** | RD | F#1 | 17 | 74732959 | SRSF2 | NM_003016 | c.284C>A | p.P95H | 43.85% | ONCOGENIC |
| **27** | RD | F#1 | X | 53430498 | SMC1A | NM_006306 | c.2420G>A | p.R807H | 10.94% | UNKNOWN |
| **28** | RD | F#1 | 1 | 115258748 | NRAS | NM_002524 | c.34G>A | p.G12S | 6.32% | ONCOGENIC |
| **28** | RD | F#1 | 4 | 55599321 | KIT | NM_000222 | c.2447A>T | p.D816V | 10.08% | ONCOGENIC |
| **28** | RD | F#1 | 17 | 74732959 | SRSF2 | NM_003016 | c.284C>A | p.P95H | 47.38% | ONCOGENIC |
| **29** | RD | F#1 | 21 | 36164485 | RUNX1 | NM_004364 | c.1390A>T | p.T464S | 9.50% | UNKNOWN |
| **30** | RD | F#1 | 4 | 106156364 | TET2 | NM_001127208 | c.1270dupA | p.S424Kfs*19 | 38.97% | Likely ONCOGENIC |
| **30** | RD | F#1 | 4 | 106182916 | TET2 | NM_001127208 | c.3957delA | p.E1320Rfs*43 | 44.09% | Likely ONCOGENIC |
| **30** | RD | F#1 | 17 | 74732959 | SRSF2 | NM_003016 | c.284C>A | p.P95H | 4.52% | ONCOGENIC |

| | | | | | | | | | | |
|---|---|---|---|---|---|---|---|---|---|---|
| **30** | RD | F#1 | X | 123197782 | STAG2 | NM_001042 749 | c.1907dupA | p.Y636* | 5.80% | Likely ONCOGENIC |
| **30** | RD | F#1 | X | 133511705 | PHF6 | NM_032458 | c.58T>A | p.C20S | 89.86% | UNKNOWN |
| **31** | RD | F#1 | 21 | 44524456 | U2AF1 | NM_006758 | c.101C>T | p.S34F | 12.34% | ONCOGENIC |
| **31** | RD | F#1 | 21 | 36252878 | RUNX1 | NM_001754 | c.484A>G | p.R162G | 7.03% | Likely ONCOGENIC |
| **31** | RD | F#1 | 21 | 36231774 | RUNX1 | NM_001754 | c.593_609delATGGGC CCCGAGAACCT | p.D198Afs*9 | 6.21% | ONCOGENIC |
| **32** | RD | F#1 | 17 | 7578404 | TP53 | NM_000546 | c.526T>G | p.C176G | 2.15% | ONCOGENIC |
| **32** | RD | F#1 | 17 | 7578406 | TP53 | NM_000546 | c.524G>A | p.R175H | 4.28% | ONCOGENIC |
| **32** | RD | F#1 | 17 | 74732959 | SRSF2 | NM_003016 | c.284C>T | p.P95L | 12.02% | Likely ONCOGENIC |
| **32** | RD | F#1 | 21 | 36231797 | RUNX1 | NM_001754 | c.587C>A | p.T196K | 16.21% | Likely ONCOGENIC |
| **32** | RD | F#1 | 18 | 42531907 | SETBP1 | NM_015559 | c.2602G>A | p.D868N | 4.74% | ONCOGENIC |
| **33** | RD | F#1 | 12 | 25398285 | KRAS | NM_033360 | c.34G>A | p.G12S | 9.41% | ONCOGENIC |
| **33** | RD | F#1 | 21 | 36231782 | RUNX1 | NM_001754 | c.602G>A | p.R201Q | 11.07% | ONCOGENIC |
| **33** | RD | F#1 | 12 | 11992079 | ETV6 | NM_001987 | c.169C>T | p.Q57* | 6.22% | UNKNOWN |
| **33** | RD | F#1 | 19 | 13054704 | CALR | NM_004343 | c.1231G>A | p.G411S | 44.64% | UNKNOWN |
| **33** | RD | F#1 | 2 | 25463286 | DNMT3 A | NM_022552 | c.2207G>A | p.R736H | 8.04% | Likely ONCOGENIC |
| **33** | RD | F#1 | 20 | 31023691 | ASXL1 | NM_015338 | c.3178dupG | p.V1060Gfs*27 | 5.66% | Likely ONCOGENIC |
| **33** | RD | F#1 | 4 | 106164741 | TET2 | NM_001127 208 | c.3609C>G | p.S1203R | 39.81% | UNKNOWN |
| **33** | RD | F#1 | 7 | 14854429 1 | EZH2 | NM_004456 | c.100C>T | p.R34* | 23.27% | UNKNOWN |
| **33** | RD | F#1 | 7 | 14854365 9 | EZH2 | NM_004456 | c.149T>C | p.L50S | 25.90% | UNKNOWN |
| **34** | IMPACT | F#1 | 21 | 36231791 | RUNX1 | NM_001754 | c.593A>G | p.D198G | 2.80% | Likely ONCOGENIC |
| **34** | IMPACT | F#1 | 17 | 74732959 | SRSF2 | NM_003016 | c.284C>A | p.P95H | 5.40% | ONCOGENIC |
| **34** | IMPACT | F#1 | 4 | 10619374 8 | TET2 | NM_001127 208 | c.4210C>T | p.R1404* | 5.30% | Likely ONCOGENIC |
| **34** | IMPACT | F#1 | 20 | 31022441 | ASXL1 | NM_015338 | c.1934dupG | p.G646Wfs*12 | 22.20% | Likely ONCOGENIC |
| **35** | IMPACT | F#1 | 21 | 36259325 | RUNX1 | NM_001754 | c.165delG | p.L56Cfs*16 | 2.00% | Likely ONCOGENIC |
| **35** | IMPACT | F#1 | 4 | 10615819 8 | TET2 | NM_001127 208 | c.2482delT | p.C828Afs*13 | 43.40% | Likely ONCOGENIC |
| **35** | IMPACT | F#1 | 4 | 10615758 0 | TET2 | NM_001127 208 | c.3100delC | p.Q1034Sfs*21 | 3.00% | Likely ONCOGENIC |
| **35** | IMPACT | F#1 | 4 | 187541413 | FAT1 | NM_005245 | c.6327C>G | p.D2109E | 9.20% | UNKNOWN |
| **35** | IMPACT | F#1 | 1 | 1:4381843 1 | MPL | NM_005373 | c.1896G>T | p.W632C | 5.50% | UNKNOWN |
| **35** | IMPACT | F#1 | 12 | 11288818 9 | PTPN11 | NM_002834 | c.205G>A | p.E69K | 12.30% | Likely ONCOGENIC |
| **35** | IMPACT | F#1 | 17 | 74732959 | SRSF2 | NM_003016 | c.284C>G | p.P95R | 13.10% | Likely ONCOGENIC |

| | | | | | | | | | | |
|---|---|---|---|---|---|---|---|---|---|---|
| **36** | IMPACT | F#1 | 20 | 31022441 | ASXL1 | NM_015338 | c.1934dupG | p.G646Wfs*12 | 17.30% | Likely ONCOGENIC |
| **37** | IMPACT | F#1 | 4 | 106196487 | TET2 | NM_001127208 | c.4823_4824delAT | p.Y1608Ffs*5 | 45.10% | Likely ONCOGENIC |
| **37** | IMPACT | F#1 | 17 | 7578208 | TP53 | NM_000546 | c.641A>G | p.H214R | 39.70% | Likely ONCOGENIC |
| **37** | IMPACT | F#1 | 17 | 74732959 | SRSF2 | NM_003016 | c.284C>A | p.P95H | 44.60% | ONCOGENIC |
| **38** | RD | F#1 | 21 | 36164448 | RUNX1 | NM_001754 | c.1427A>C | p.V476A | 1.20% | UNKNOWN |
| **38** | RD | F#1 | 1 | 36932040 | CSF3R | NM_000760 | c.2429A>G | p.D810G | 46.10% | UNKNOWN |
| **38** | RD | F#1 | 5 | 170837543 | NPMI | NM_002520 | c.860_863dupTCTG | p.W288Cfs*12 | 3.30% | Likely ONCOGENIC |
| **38** | RD | F#1 | 2 | 209113113 | IDH1 | NM_005896 | c.394C>A | p.R132S | 2.40% | Likely ONCOGENIC |
| **39** | IMPACT | F#1 | 6 | 15374449 | JARID2 | NM_004973 | c.148_149insCG | E50Afs*8 | 87.00% | Likely ONCOGENIC |

RD, raindance; D, diagnosis; F, follow-up.

Supplemental Table 6 Summary of primary PDX mice

| Patients | NSG mice | Cells injected/mouse | hCD45 (%) in PB | | hCD45 (%) in BM | Leukemic blasts | pDCs | Lymphocytes | Transplantable in 2° PDX mice |
|---|---|---|---|---|---|---|---|---|---|
| | | | 3 months | 6 months | | | | | |
| **RC** | 1 | 1.3 million | 2.86 | 59.2 | 99 | Y | Y | N | Y |
| | 2 | 1.3 million | 4.44 | 55.9 | 95 | Y | Y | N | Y |
| | 3 | 1.3 million | 7.28 | 53.7 | 97.1 | Y | Y | N | Y |
| **ES** | 1 | 374 000 | 15.3 | 16 | 5 | N | N | Y | N |
| | 2 | 374 000 | 43.9 | | 35 | N | N | Y | N |
| **RM** | 1 | 300 000 | 0.39 | 2.59 | 0.9 | N | N | Y | N |
| | 2 | 300 000 | 0.36 | 5.98 | 1.0 | N | N | Y | N |
| **GR** | 1 | 186 000 | | 1.85 | 0.5 | N | N | Y | N |
| | 2 | 186 000 | | 1.80 | 0.4 | N | N | Y | N |
| **DA** | 1 | 600 000 | 0.15 | | 0.09 | N | N | Y | N |
| | 2 | 600 000 | 0.16 | | 0.2 | N | N | Y | N |
| | 3 | 600 000 | 0.18 | | 0.1 | N | N | Y | N |
| **RG** | 1 | 2 million | 10 | | 4.22 | N | Y (0.2%) | Y | N |
| | 2 | 2 million | 5.1[a] | | | | | | |
| | 3 | 2 million | 24.5 | | 7.64 | N | Y (1.8%) | Y | N |
| | 4 | 2 million | 70 | | 18.9 | N | Y (2.6%) | Y | N |
| | 5 | 2 million | 10.3[a] | | | | | | |
| | 6 | 2 million | 8.3[a] | | | | | | |
| | 7 | 2 million | died | | | | | | |

a: mice died after 3 months evaluation.
